# Host transcriptional responses to gut microbiome variation arising from urbanism

**DOI:** 10.1101/2025.10.26.683539

**Authors:** Sabrina Arif, Shreya Nirmalan, Adnan Alazizi, Henriette Mair-Meijers, Adwoa Agyei, Mary Y. Afihene, Shadrack O. Asibey, Yaw A. Awuku, Amoako Duah, Amelie Plymoth, Yvonne Nartey, Lewis Roberts, Kenneth Valles, Fatimah Ibrahim, Yvonne A. L. Lim, Tan Maw Pin, Charles Onyekwere, John Rusine, Ivan E Mwikarago, Eric J. Alm, Mathilde Poyet, Mathieu Groussin, Francesca Luca, Ran Blekhman

**Affiliations:** Section of Genetic Medicine, Department of Medicine, University of Chicago, Chicago, IL, USA; Center for Molecular Medicine and Genetics, Wayne State University, Detroit, Michigan 48201, USA; Global Microbiome Conservancy; Department of Biological Engineering, Massachusetts Institute of Technology, Cambridge, MA, United States; Institute of Experimental Medicine, Kiel University, Kiel, Germany; Institute of Clinical Molecular Biology, Kiel University, Kiel, Germany; Department of Human Genetics, University of Chicago, Chicago, IL, USA; Department of Medicine and Therapeutics, University of Ghana Medical School and Korle Bu Teaching Hospital, Accra, Ghana; Department of Medicine, Kwame Nkrumah University of Science and Technology, Kumasi, Ghana; Catholic University College, Sunyani, Ghana; Department of Internal Medicine and Therapeutics, School of Medical Sciences University of Cape Coast, Cape Coast, Ghana; Department of internal medicine, University of Ghana Medical Centre, Legon, Accra, Ghana; Department of Medical Epidemiology and Biostatistics, Karolinska Institutet, Stockholm, Sweden; Division of Gastroenterology and Hepatology, Mayo Clinic, 200 First Street SW, Rochester, MN, USA; Medical Scientist Training Program, Mayo Clinic, Rochester, United States; Centre for Innovation in Medical Engineering, University of Malaya, Kuala Lumpur, Malaysia; Department of Parasitology, Faculty of Medicine, Universiti Malaya, Kuala Lumpur, Malaysia; Faculty of Medicine, University of Malaya, Kuala Lumpur, Malaysia; Department of Medicine, Lagos State University College of Medicine, Lagos, Nigeria; National Reference Laboratory, Kigali, Rwanda; National Institute of Allergy and Infectious Diseases (NIAID), Rockville, Maryland, United States; Rwanda Food and Drug Authority, Kigali, Rwanda, and College of Medicine and Health Sciences, University of Rwanda; The Broad Institute of MIT and Harvard, Cambridge, MA, United States

**Author notes:** Corresponding authors: Sabrina Arif Ran Blekhman, Francesca Luca. These authors contributed equally to this work (Adwoa Agyei, Mary Y. Afihene, Shadrack O. Asibey, Yaw A. Awuku, Amoako Duah, Amelie Plymoth, Yvonne Nartey, Lewis Roberts, Kenneth Valles, Fatimah Ibrahim, Yvonne A. L. Lim, Tan Maw Pin, Charles Onyekwere, John Rusine, Ivan E Mwikarago, Eric J. Alm). These authors contributed equally to this work (Francesca Luca, Ran Blekhman). microbiomeconservancy.org.

## Abstract

Gut microbiomes of urban communities are compositionally different from their rural counterparts, and are associated with immune dysregulation and gastrointestinal disease. However, it is unknown whether these compositional differences impact host physiology, and through what mechanisms. Here, we used human colonic epithelial cells to directly compare host transcriptional changes induced by gut microbiomes from urban versus rural communities. We co-cultured host cells with live, stool-derived gut microbiomes from Rwanda, Ghana, Nigeria, Malaysia, and the United States, and quantified transcriptional responses using RNA-seq. We found that urban microbiomes affected innate immune pathways, including TNF signaling and bacterial antigen recognition. We also found that high-diversity microbiomes elicited a stronger host transcriptional response, while low-diversity microbiomes triggered epithelial restructuring and glycolysis. Finally, specific taxa driving these effects, including *Bifidobacterium adolescentis* and *Bacteroides dorei*, correlated with lifestyle factors such as diet. These findings demonstrate that urbanization-associated microbiome changes directly influence host epithelial gene expression.

## INTRODUCTION

Variation in the composition of the gut microbiome—the community of microorganisms inhabiting the human intestine—is linked to a wide range of health outcomes, motivating studies of lifestyle factors that influence gut microbiome composition [1–3]. Globally, a prominent source of gut microbiome variation is industrialization and urbanism [4–8]. Industrialization describes broad socioeconomic and lifestyle transitions associated with industrial economies, while urbanism reflects more localized differences along the rural-to-urban gradient. Although not identical, both processes are characterized by high sanitation, medication use, and intake of processed foods. Gut microbiomes from urban communities (urban gut microbiomes) are typically dominated by the genus *Bacteroides*, *Bifidobacterium*, *Ruminococcus*, and by microbial functions suited for simple sugar metabolism and lipid and fat metabolism. In contrast, rural gut microbiomes are enriched for several species of *Prevotella*, Spirochaetes and other organisms known for complex carbohydrate metabolism and production of beneficial short chain acids [6,9].

Non-communicable intestinal diseases characterized by chronic inflammation are rising globally alongside urbanism and industrialization, often outpacing the local health system’s capabilities to effectively care for patients [10–13]. Because individuals leading urban lifestyles have altered microbiome composition relative to those leading rural lifestyles, several studies hypothesized that the microbiome mediates the association between non-communicable intestinal diseases and industrialization [4,14–17]. Additionally, observational surveys and diet interventions have identified correlations between gut microorganisms enriched in urban microbiomes and inflammatory response [18–21], however, limited incorporation of gut microbiomes from rural cohorts constrains interpretations. Moreover, the mechanism through which the microbiome interacts with the host to promote disease phenotypes associated with urbanism and industrialization remains unclear.

One hypothesis is that gut microbiome taxa influence the expression of genes regulating inflammatory pathways linked to disease. It is well established that the microbiome modulates host gene expression in the interfacing epithelium [22,23] and recent studies have observed disease-specific associations between the microbiome and host gene expression [24–26]. Experimentally validating these microbiome–host interactions through a combination of metagenomic shotgun sequencing of microbial communities and RNA sequencing of the interfacing gut epithelial cells may yield important insights into the relationship between microbiome composition and gut health. Human intestinal epithelial cell cultures have become valuable tools for investigating the mechanisms of gene expression of the intestinal epithelium *in vitro* and offer a complementary approach to clinical and animal studies, while allowing for tightly-controlled, high-throughout, and cost-effective experimental designs [27,28]. We previously established a colonic epithelial cell culture system to quantify host gene expression response to live microbial communities. We used this experimental system to study host gene expression changes associated with microbiome variation between individuals and primate species [29–31]. However, the transcriptional response of host epithelium to variation in the microbiome arising from urbanism remains unexplored.

Here, we assessed how host gene regulation is impacted by taxonomic and functional variation in gut microbiome composition that is associated with urbanism. Using microbiome samples from individuals leading urban and rural lifestyles, we used a combination of host mRNA sequencing and microbiome metagenomic shotgun sequencing to determine host genes and pathways affected by urban compared to rural microbiomes, and evaluated the contributions to the host response from individual microbial species and microbial functions. This experimental design allowed us to directly test the effects of lifestyle associated microbiome variation on host cells while controlling for technical and environmental factors, shedding light on the contributions of the gut microbiome to host physiology.

## RESULTS

To quantify how variation in microbiome composition arising from urbanism impacts gene expression in host cells, we extracted live microbiota from stool samples of individuals leading urban and rural lifestyles across Ghana, Nigeria, Rwanda, Malaysia, and the United States (39 urban and 24 rural, **Table S1**) and characterized their taxonomic composition. We treated human colonic epithelial cells (colonocytes) with these live microbiomes using an experimental system that has been previously utilized by our laboratory [29–31] and measured the host transcriptional response using RNA-seq (**Figure 1A**).

**FIGURE 1.**
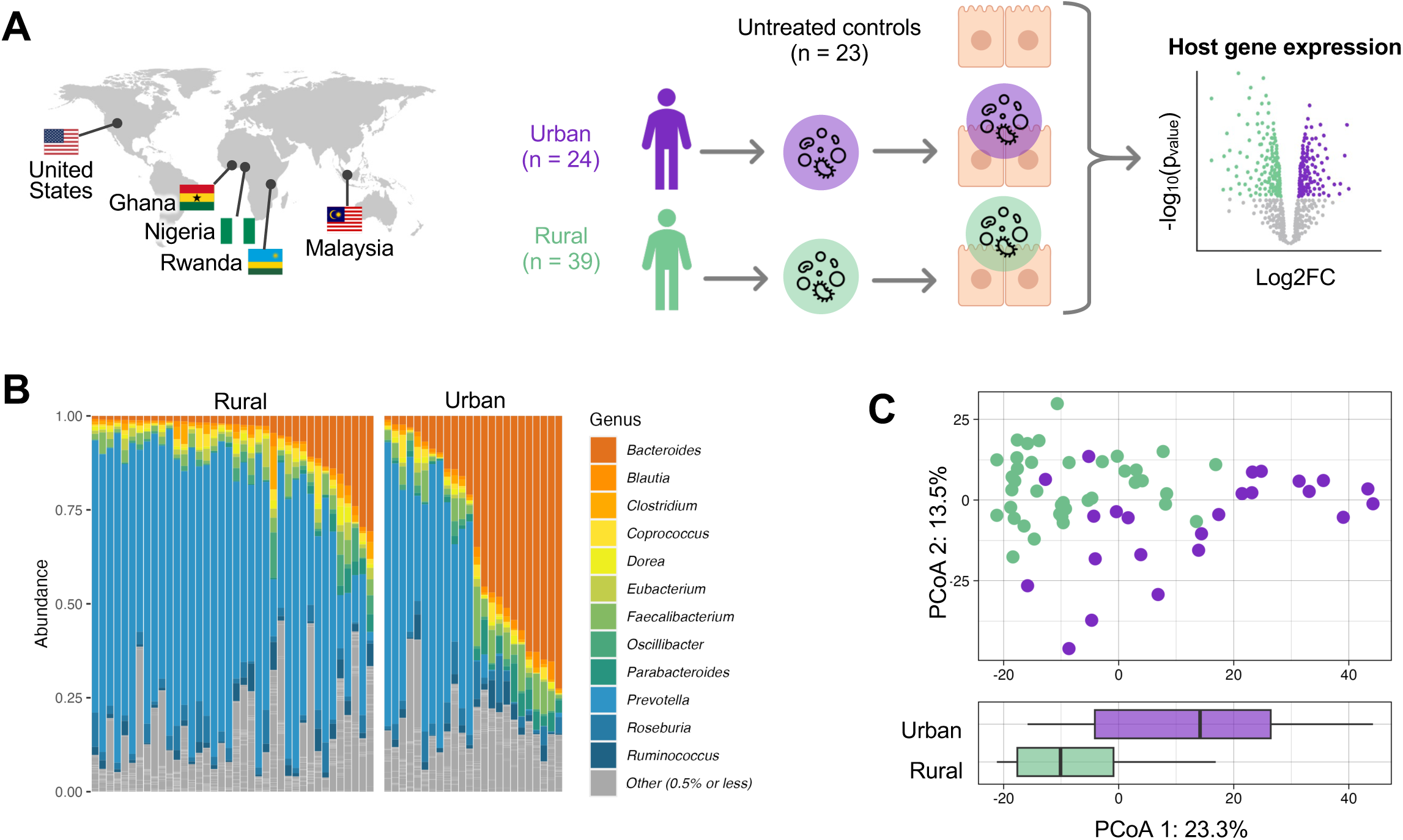
Experimental model and microbiome sample set overview. **A** Fecal microbiome samples of urban (n=24) and rural (n=39) communities were sampled from five countries. Colonocytes were treated (n=63; n=23 control) with the live microbiome samples (see Methods) and gene expression was measured using RNA sequencing. **B** The y-axis represents genus-level relative abundances for the microbiomes of 63 donors (x-axis). Taxa in color have a median abundance of 0.5% or greater while low-abundance genera are grouped together in grey (see legend). **C** Top: Genus-level principal coordinate analysis (PCoA) of microbiome samples. The x-and y-axes are the first and second principal components, representing 23.3% and 13.5% of variability in microbiome composition, respectively. Bottom: Box plot of PC1. Values of PC1 are shown for urban (purple) and rural (green) samples.

First, we assessed the taxonomic and functional composition of the microbiomes via shotgun metagenomic sequencing. We found that microbiome composition reflects known patterns reported in previous surveys of urban and rural cohorts [4,9] (**Figure 1B**). We identified 9 families, 11 genera, and 33 species that were differentially abundant between urban and rural microbiomes (FDR-corrected *p*-value < 10% using ALDEx2; **Methods**). *Bifidobacterium, Alistipes*, and *Akkermansia*, along with several species of *Bacteroides*, characterized urban microbiomes at the genus level. These genera are commonly found in surveys of urban communities, with *Alistipes* and *Bacteroides* notable for their implications in numerous conditions, including colorectal cancer and inflammatory bowel disease [26,32,33]. In contrast, we found that rural microbiomes had a higher abundance of several species of *Prevotella*, many of which are uncharacterized (**Figure S2**). *Prevotella* is broadly linked to fiber-rich diets, especially in rural communities [34,35]. In fact, these specific novel species have been previously identified in the gut microbiomes of the Hadza, a nomadic population in rural Tanzania that subsists primarily on hunted and foraged foods [6,35]. Analysis in a larger cohort showed that many of these taxa remain significantly associated with industrialization and urbanism even after accounting for diet, genetics, geography, age, sex, and BMI [8]. Urbanism accounted for much of the variation among genus-level microbiome composition (**Figure 1C**, *p*-value = 7.15×10^-7^; Wilcoxon rank sum exact test). Overall, these results indicate that the gut microbiomes used in our study recapitulate known patterns in urban and rural communities.

### Urban microbiomes influence innate immune pathways in the host

To investigate whether microbiome features arising from urbanism are associated with changes in host gene expression, we performed RNA-seq in colonocytes treated with either urban or rural microbiomes and untreated controls (n=24, 39, and 23, respectively, **Figure S3**, **Methods**). We found that, relative to the control, the urban microbiomes elicited a stronger transcriptional response in the colonocytes compared to rural microbiomes, as indicated by greater absolute effect sizes (*p*-value = 1.30×10^-3^; exact binomial test; **Figure 2A, Figure S4**). Of the host genes that responded to the live microbiomes, 663 were significantly differentially expressed in response to urban microbiomes, and 472 to rural microbiomes (FDR < 10% using DESeq2; **Methods**). Among these, 292 host genes responded to both urban and rural microbiomes, while 371 and 180 host genes were significant only for the urban and rural microbiome conditions, respectively.

**FIGURE 2.**
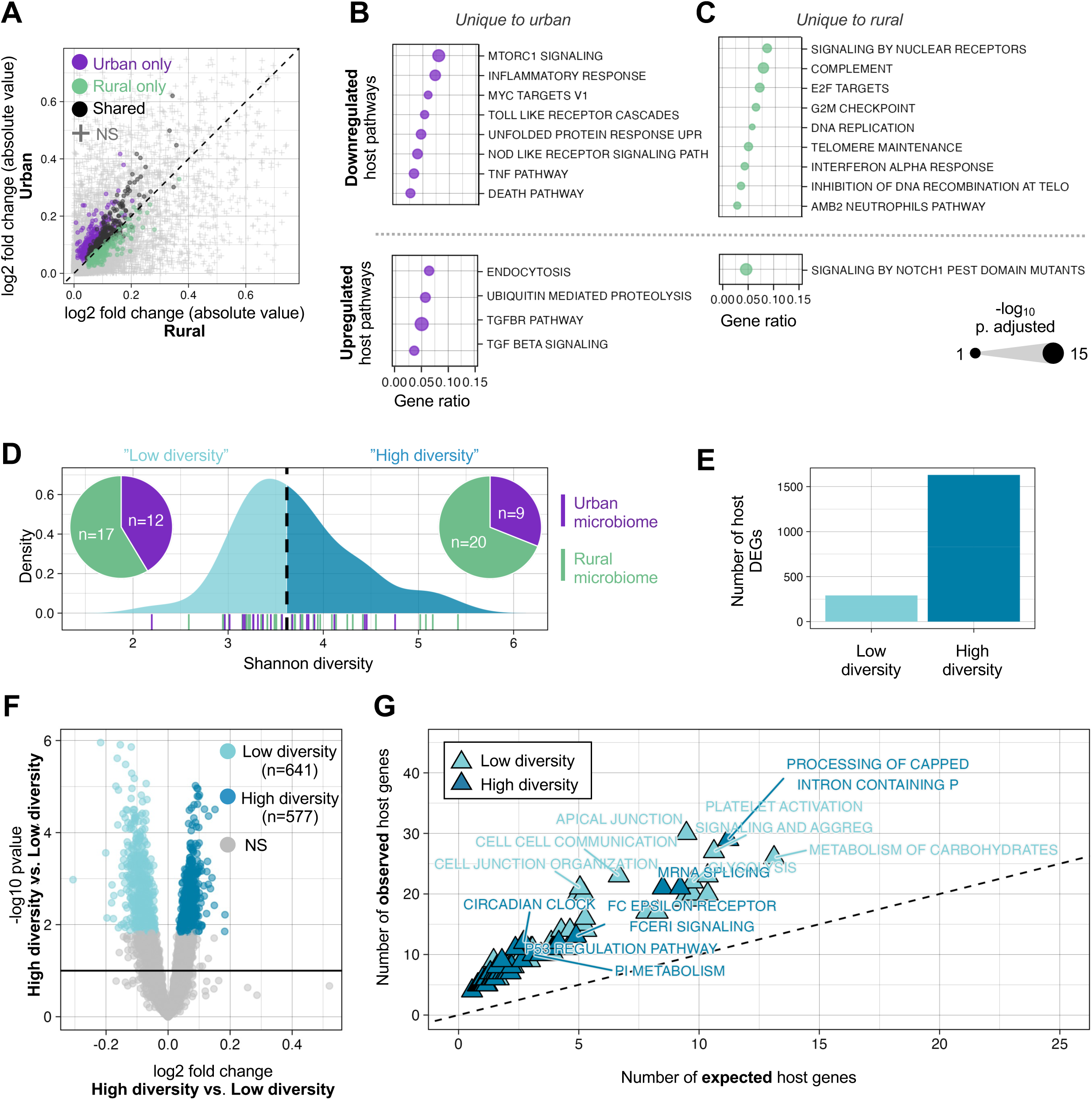
A–C Host transcriptional response to urban and rural microbiomes. **A** Differentially expressed host genes (DEGs) compared to control condition. Log₂ fold change values shown for DEGs significant at FDR < 10%. Dots indicate DEGs specific to urban (purple), rural (green), or shared between both (black) microbiome conditions. **B, C** Enriched host gene pathways (FDR < 10%, gene count ≥ 3) in response to urban (B) and rural (C) microbiomes. Upregulated and downregulated pathways shown in bottom and top panels, respectively. Dot size reflects –log(p-value). **D–G Microbiome alpha diversity associated with distinct host transcriptional responses. D** Density plot of Shannon diversity values across microbiome samples. Median (3.62) indicated by black dotted line. Samples below the median (light blue) are classified as “low diversity”; those above (dark blue) as “high diversity.” Pie charts indicate the proportion of urban (purple) and rural (green) samples in each group. Rug plot shows individual sample distribution: purple and green ticks correspond to urban and rural communities, respectively. **E** Number of host DEGs in response to low diversity microbiomes vs. control (light blue) and high diversity microbiomes vs. control (dark blue). **F** Volcano plot depicting host genes that respond to low diversity vs. high diversity microbiomes. Genes significantly (FDR < 10%) upregulated in low diversity microbiomes shown in light blue, high diversity shown in dark blue, and NS shown in grey. **G** Enriched host gene pathways (FDR < 10%, gene count ≥ 3) in response to low (light blue) vs. high (dark blue) diversity microbiomes. Observed values (y-axis) represent the number of input genes annotated to each pathway. Expected values (x-axis) were calculated as the input list size multiplied by the proportion of background genes annotated to the pathway; 1:1 shown by black dashed line.

We next performed overrepresentation analysis on each condition separately to provide functional context to the host genes modulated by urban and rural microbiomes. We found that host genes downregulated in response to both urban and rural microbiomes were enriched in pathways involved in innate immune response (TNF signaling via NF-κB, IL10 signaling). Conversely, host genes upregulated in response to both urban and rural microbiomes were enriched in functions related to hypoxia and UV response (**Figure S4**). We also found that host genes that responded uniquely to urban or rural microbiomes were enriched in specific pathways. For example, host genes downregulated only in response to urban microbiomes were enriched in bacterial antigen recognition and cell turnover/barrier function (Toll-like receptor cascades, NOD-like receptor signaling and MTORC1 signaling respectively). TGF-β signaling, a host pathway known for its immuno-suppressive properties [36], was enriched in response only to urban microbiomes (**Figure 2B**). In contrast, several of the host pathways enriched in response to only rural microbiomes, such as E2F targets and telomere maintenance, were related to DNA replication and damage repair (**Figure 2C**).

### Microbiome diversity drives host immune and epithelial integrity pathways

Previous surveys have reported increased taxonomic diversity in rural microbiomes compared to those from urban communities [4,6,17]. This trend is also confirmed in the broader GMbC cohort [8], and a tendency for higher diversity in rural communities is observed in our dataset (**Figure S5**). Thus, we next questioned whether microbiome diversity influenced host gene expression responses. To answer this question, we quantified alpha diversity in all microbiome treatments and split our samples into low-diversity and high-diversity subsets of equal size (**Figure 2D**; see Methods). Relative to controls, low-diversity microbiomes induced significant changes in expression for 224 host genes, while high-diversity microbiomes induced changes in 1,386 host genes (**Figure 2E**; FDR < 10%).

We next directly compared host responses to low diversity and high diversity microbiomes, independent of controls. Host genes expressed in response to low-diversity communities were enriched in pathways supporting epithelial integrity and barrier function, including cell junction and apical junction organization, keratinization, and epithelial–mesenchymal transition (**Figure 2F, Table S5**). These processes are involved in maintaining epithelial integrity; for instance, apical junction components regulate tight and adherens junctions, which control cellular permeability and protect against microbial translocation in the gut [37,38]. In addition, functions related to cellular energy metabolism such as glycolysis and carbohydrate metabolism were upregulated in response to low-diversity microbiomes. In contrast, high diversity microbiomes promoted expression of pathways related to broad regulatory networks. This included pre and post-translational modification (mRNA splicing, processing of capped introns in pre-mRNA, neddylation), and receptor-mediated signaling (ERBB1 and FCER1 signaling) (**Figure 2G, Table S5**).

### Host gene response to abundance of individual taxa show lifestyle-specific patterns

Although our initial analysis revealed broad changes in host gene expression in response to microbiomes from urban and rural communities, it did not resolve which specific microbial taxa are driving these changes. Thus, we investigated how expression of individual host genes changed in response to the abundance of individual taxa in our study system using a lasso penalized regression model to identify host gene-microbe associations (FDR < 10%; **Methods**). In urban microbiomes, we observed a total of 347 distinct host gene–microbe associations, with 84 unique taxa driving the expression of these host genes. Colonocytes treated with rural microbiomes displayed 610 total significant associations comprising 126 taxa and 575 host genes (**Figure 3A**). Across both urban and rural communities, a few taxa consistently drove the expression of many host genes. For example, in urban microbiomes, the abundance of *Dorea A longicatena B* impacted the expression of 12 host genes, while in rural microbiomes this microbe drove expression of 30 host genes. *RUG115 sp900066395* was another influential taxon, which led to the change in the expression of 10 and 24 host genes in urban and rural microbiomes, respectively. Overall, the microbes that showed the largest impact on host gene expression included well-known short chain fatty acid producers belonging to the family Lachnospiraceae, such as *Dorea*, *Roseburia*, *Coprococcus*, as well as the genera *Clostridium* and *Ruminococcus* (**Figure 3B, Table S2**). Lachnospiraceae is a family of fermentative and anaerobic organisms that encodes numerous beneficial functions for the host and is found in increased abundance near gut mucosa *in vivo* [39,40].

**FIGURE 3.**
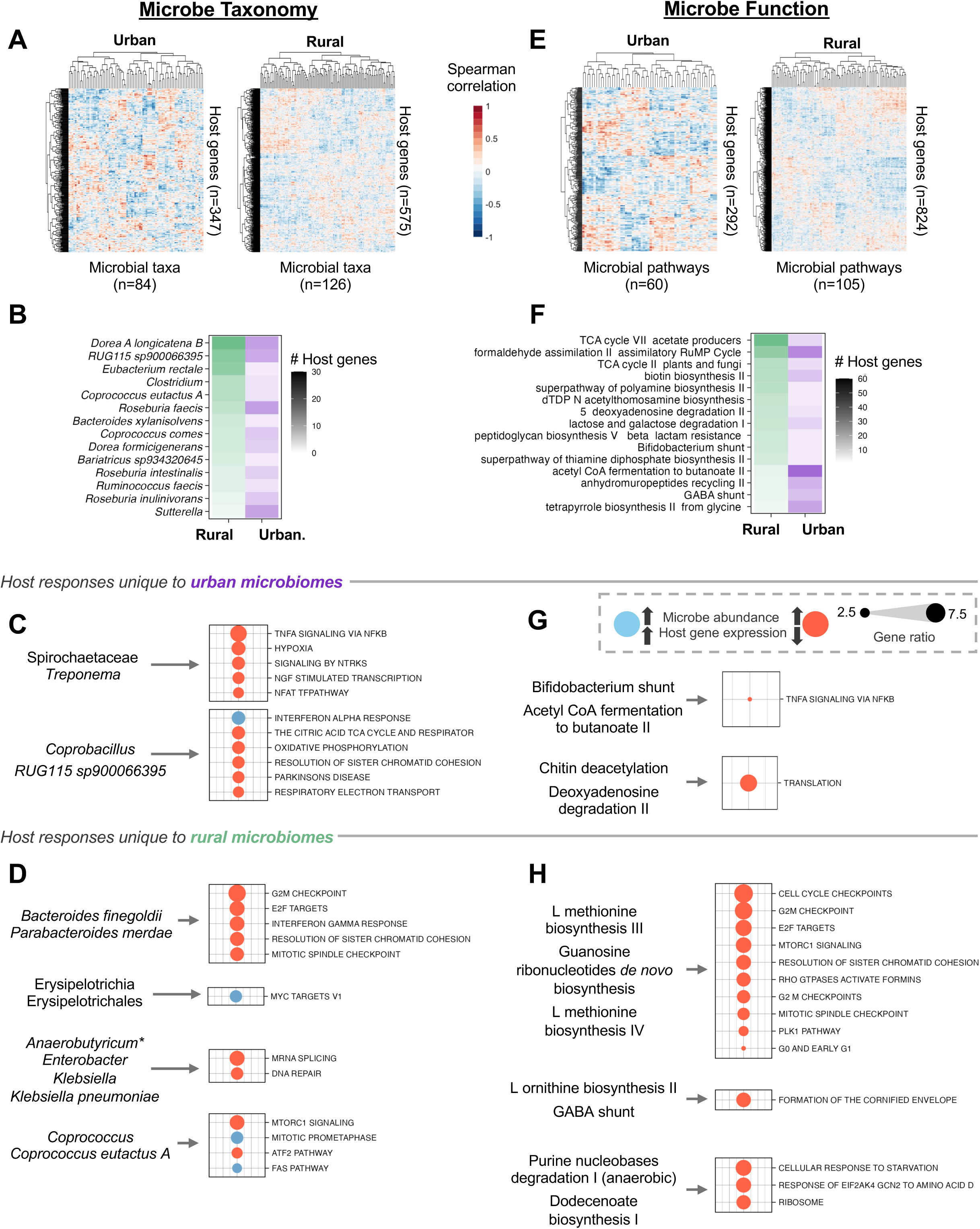
A-D Transcriptional response of host to microbe taxonomy. **A** Spearman correlation between microbial taxa (x axis) and expression levels of host genes (y axis). Value of Spearman correlation coefficient indicated by color (red=positive, white=0, blue=negative). Taxa/host gene only included if identified as significant (FDR < 10%) in lasso analysis (see Methods). Analysis applied separately to colonocytes responding to urban (left) and rural (right) microbiomes. x and y axes subjected to hierarchical clustering using the Euclidean distance metric. **B.** Microbes (y axis) with the greatest number of pairwise host gene–microbe associations in lasso analysis. Number of host gene–microbe associations depicted by darkness of shading; green and purple shading reflect effect in rural and urban microbiomes, respectively (x axis). **C, D** Associative clusters between groups of taxa and groups of host genes in urban (C) and rural (D) samples Left text indicates groups of microbe taxa identified by canonical correlation analysis as having associations with groups of host genes. We only show associative clusters with significant over-representation. Right text indicates significant (FDR < 10%, gene count ≥ 3) host gene pathways enriched in each associative cluster. Dot size represents gene ratio: the proportion of input host genes that are annotated in a term. Blue dots indicate positive associations, orange dots indicate negative associations. Unless indicated by an asterisk (*Anaerobutyricum*), all microbes in a cluster act in the same direction. **E-H Transcriptional response of host to microbe functions. E** Similar to A, but with microbial pathways on the x axis. **F** Similar to B, but with microbial pathways on the y axis. **G, H** Similar to C, D, but considering microbial pathways. Groups of microbial pathways (left) and significant associated host gene pathways as identified by overrepresentation analysis (right).

Some instances of host gene expression responses to specific microbes appeared to be dependent on the overall microbiome context: 25 species drove host gene expression only when present in urban microbiomes, and 67 species drove host gene expression only when present in rural samples (**Table S2**). In rural microbiomes, over 30% of these 67 species drove expression of more than three host genes, compared to less than 10% of species in urban microbiomes. For example, *Eubacterium rectale*, a ubiquitous organism capable of complex carbohydrate utilization and short-chain fatty acid production [41], influenced the expression of 22 host genes in rural microbiomes, but zero in urban microbiomes. The same was true of a novel species of *Akkermansia*, *sp004167605*, which drove the expression of 14 host genes in rural microbiomes and none in urban microbiomes. *Akkermansia* is known for its positive roles in modulating host metabolism in health and antagonistic roles in disease [42]. Conversely, members of the genus *Sutterella* were more influential in urban microbiomes. Specifically, *Sutterella wadsworthensis*, and *Sutterella* at the genus level drove the expression of more host genes in urban microbiomes compared to rural microbiomes (19 and 11, respectively and 4 and 0, respectively; **Figure 3B**). *Sutterella* species, historically regarded as commensals with mild pro-inflammatory characteristics, have gained recent interest for their enrichment in obesity and ulcerative colitis cohorts [43].

Interestingly, many taxa that were differentially abundant among urban and rural microbiomes had minimal impacts on host gene expression. For example, several species that were significantly more abundant in urban communities, such as *Bacteroides vulgatus, Bacteroides caccae, Bacteroides ovatus,* and *Alistipes putredinis*, affected zero host genes both when in the context of urban or rural microbiomes. Likewise, species of *Prevotella,* a hallmark of rural microbiomes, also lacked any influence on host gene expression (**Figure S2, Figure S6**). However, we note that some taxa that impacted host gene expression in this dataset, such as *Eubacterium* and members of the family Clostridiaceae, are typically found in rural microbiomes in larger studies [4,44].

While prior studies support the existence of interactions in which the abundance of a specific taxon drives the expression of an individual host gene [45], another possibility is that groups of microbes with correlated abundance patterns influence the concerted expression of multiple host genes [24]. To address this possibility, we utilized canonical correlation analysis to identify host gene–microbe “clusters”, or correlations between groups of microbes and groups of host genes (**Table S4**, **Methods**). To provide functional context to the host genes found in each cluster, we performed overrepresentation analysis. In host cells treated with urban microbiomes, the expression of host genes involved in cellular immune modulation by bacteria, such as hypoxia, and TNF signaling [46,47], decreased with the increased abundance of the family Spirochaetaceae and the genus *Treponema* (**Figure 3C**). Interestingly, *Treponema* has also been identified as negatively associated with calprotectin levels, a marker of gut inflammation, suggesting its potential role as a marker of healthy microbiomes [8]. We also found the expression of genes in the interferon alpha response to cellular respiration (oxidative phosphorylation, respiratory electron transport) was mediated by *Coprobacillus* and *RUG115 sp900066395* in urban microbiomes. In host cells treated with rural microbiomes, we observed attenuated expression of host genes in pathways related to DNA replication and repair, such as G2M checkpoint and E2F targets, with higher abundance of *Bacteroides finegoldii* and *Parabacteroides merdae* (**Figure 3D**). Genes involved in processes related to programmed cell death, such as the Fas pathway, were upregulated with higher abundance of *Coprococcus*, while those in pathways involved in mTOR signaling had lower expression.

### Host genes responses to microbial functions are lifestyle specific

Microbial functions, rather than individual taxa, may be the most critical drivers of host– microbiome interactions, as taxonomic composition alone does not always reflect what microbes are doing; closely related microbes can perform different functions, and similar functions can arise across diverse taxa [48,49]. Thus, to determine how microbial function influences host gene expression, we used lasso penalized regression (FDR < 10%; **Methods**) to identify individual host genes associated with microbial pathways. In samples treated with urban microbiomes, 292 host genes responded to 60 microbial pathways, whereas in rural microbiomes, 824 host genes responded to 105 pathways (**Figure 3E**). These associations pointed to distinct microbial functions shaping host responses across conditions. For instance, the pathway acetyl-CoA fermentation to butanoate II, which is central to butyrate production, a key energy source for colonic epithelial cells [50,51], was strongly linked to gene expression in urban microbiomes (41 host genes), compared to only 6 associations in the rural microbiomes. In contrast, rural microbiomes exhibited stronger associations with energy metabolism pathways: TCA cycle VII and TCA cycle II were linked to 60 and 27 host genes, respectively, compared to 7 and 5 in urban microbiomes. Additionally, the GABA shunt, a succinate-producing pathway that feeds into the TCA cycle [52,53], influenced expression of 14 host genes specifically in the rural condition (**Figure 3F**).

Similar to our analysis of host genes and microbial taxa, we used canonical correlation analysis to identify groups of host genes whose expression was driven by groups of microbial pathways (**Table S4**, **Methods**). In urban microbiomes, the abundances of the microbial pathways of the *Bifidobacterium* (Bifid) shunt and acetyl coA fermentation to butanoate II downregulated the expression of 221 host genes, which were implicated in TNFα signaling via NF-κB in the host (**Figure 3G**). Both of these microbial pathways are carbohydrate utilization pathways that are responsible for the production of short chain fatty acids, notably acetate and lactate in the Bifid shunt and acetate in acetyl coA fermentation [54,55]. Further, the Bifid shunt has previously been implicated in protection against infection by epithelial cells [56,57]. In rural microbiomes, the microbial pathways of L-methionine biosynthesis III and IV and guanosine ribonucleotides *de novo* biosynthesis decreased expression of 233 host genes (**Figure 3H**). These genes were enriched in several cell cycle-related host pathways, including G2M checkpoint, E2F targets, and RHO GTPases activate formins. L-ornithine biosynthesis II and GABA shunt drove host processes related to keratinization (i.e., formation of the cornified envelope).

### Host lifestyle influences abundance of microbes that regulate host gene expression

The descriptor “urbanism” reflects population density estimates from the SEDAC Population Estimation Service (see **Methods**) and represents broad differences in sanitation, environmental exposure, infrastructure, lifestyle, and other characteristics that lead to changes in microbiome composition [8]. Having examined this broad category, we next questioned how these individual lifestyle traits and resultant host physiological traits (referred to as host features) correlated with the abundance of organisms that drove host gene expression. These features, aggregated from survey questionnaire data and biomarker assays, included medication use, stool biomarkers, and other features such as weekly exercise and population density (**Figure S7**). Similarly, we investigated correlations between the frequency of consumption of specific foods (“diet features”) and microbial abundance. Using these diet features, we generated a “Diet PC1” metric to capture overall dietary variation across samples (**Methods**). Higher Diet PC1 values generally indicated dietary habits more characteristic of urban lifestyles, such as higher intake of processed foods and animal fat (**Figure S7**).

We then tested how host features are associated with taxa that significantly influenced host gene expression, and found 104 associations between 14 host features and 69 taxa (adjusted *p-*value < 10% generalized linear model (ALDEx2); see Methods; **Figure 4A**, **Table S2**). For example, high consumption of lactose and vegetables correlated with the abundance of *Bifidobacterium adolescentis* (Holm-corrected *p* = 6.06×10^-2^ and 8.18×10^-2^ respectively by Wald test; **Figure 4B, Figure S8**). This highly prevalent species is known for its fermentation of plant-derived glycans and production of GABA [53,58,59]. *TKT*, a crucial host gene encoding an enzyme (transketolase) that maintains ATP production in gut epithelium and an important regulator of cancer [60,61], responded to *B. adolescentis*. *Bacteroides dorei*, a species found more abundantly in urban microbiomes, correlated with Diet PC1 and increased Immunoglobulin A (IgA) in the stool (Hom-corrected *p* = 3.58×10^-3^ and 1.41×10^-2^ respectively by Wald test; **Figure 4C, Figure S2, Figure S8**). In response, the host expressed *CCN1*, which encodes a critical protein involved in inflammation and bacterial clearance and is linked to chronic intestinal diseases and cancer [62,63]. Finally, an uncharacterized species of *Phascolarctobacterium* had a negative association with Diet PC1 (Holm-corrected *p* = 3.45×10^-3^; Wald test; **Figure 4D, Figure S8**). *Phascolarctobacterium* is an asaccharolytic organism that synthesizes acetate and propionate [64,65] and impacted expression of over 30 host genes, which are enriched for the retinoic acid signaling pathway. This host pathway governs gut epithelium homeostasis and has previously been shown to be modulated by gut microbiota [66]. Eph signaling, another moderator of gut homeostasis with implications in inflammatory bowel disease [67,68], was also enriched.

**FIGURE 4.**
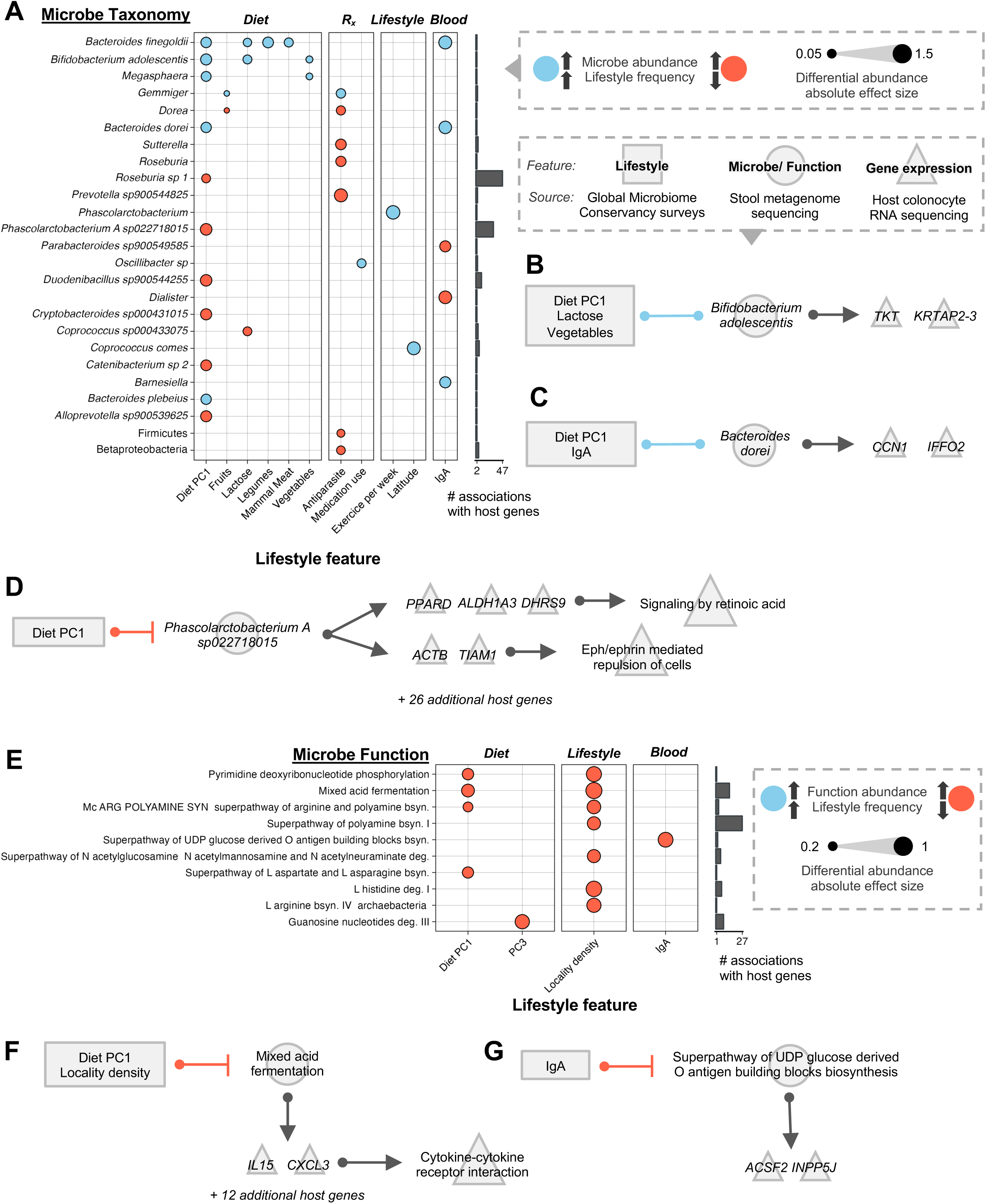
A-D Associations between host lifestyle, microbe abundance, and host gene expression. **A** Left: Association of lifestyle features (x axis left; grouped by diet, medication use, general lifestyle, and blood markers) with the abundance of microbes (y axis). Far right: number of host genes (x axis) that associate with each microbe (y axis) per lasso analysis. For simplicity, microbes are only shown if >= 1 association with a host lifestyle feature and >=2 associations with host genes. Circle indicates a significant (FDR < 10%) association between microbe and lifestyle. Dot size corresponds to absolute effect size of coefficient. Blue dots represent positive associations, orange dots represent negative associations. **B, C, D** Rectangles represent lifestyle features collected from survey data of stool donor participants. Circles represent microbe taxonomy. Triangles represent individual host genes and host pathways that are enriched from host genes. Lines between rectangles and circles represent significant (Holm-corrected *p* < 1×10^-1^) associations via differential abundance testing. Blue lines indicate positive associations, orange lines indicate negative associations. Arrows between circles and triangles represent significant (FDR < 10%) associations via lasso analysis. **E-G Associations between host lifestyle, microbe function, and host gene expression. E** Similar to A; microbe functions instead of microbe taxonomy on y axis. **F, G** similar to B, C, D; circles represent microbe function instead of microbe taxonomy.

Applying the same approach to microbial pathway abundance data, we observed 13 significant associations between 4 host features and 10 microbial functions (**Figure 4E**; see Methods and **Table S3)**. Diet PC1 and locality density were found to negatively associate with the microbial pathway of mixed acid fermentation (Holm-corrected *p* = 2.49×10^-3^ and 3.46×10^-2^, respectively; Wald test). This anaerobic process converts glycolysis end products into compounds such as succinate, 2-oxoglutarate, acetate, ethanol, and lactate [69]. Notably, this process drove the expression of two signaling molecules in the host, *IL-15* and *CXCL3*, which were broadly implicated in the host pathway of cytokine-cytokine receptor interaction (**Figure 4F**). These molecules drive pro-inflammatory processes in the host, including immune cell activation and recruitment [70–72]. Host levels of IgA correlated with a chimeric pathway implicated in the biosynthesis of O-antigens on bacterial surfaces (**Figure 4G**). IgA plays a key role in mucosal immunity by binding to surface structures such as O-antigens to neutralize antigens [73,74].

## DISCUSSION

Variation in gut microbiome diversity is a hallmark of industrialization, with industrialized and urban communities often exhibiting reduced microbial richness [3,4,6,75]. Although industrialization is frequently correlated with health outcomes, the functional consequences of microbial diversity on the host epithelium remain poorly understood. Here, we used a colonic epithelial cell culture model to test how gut microbiomes from urban and rural communities affect host gene expression. We observed more interactions with the host immune system in response to microbiomes associated with urban lifestyles, suggesting that these communities may contribute to altered inflammatory signaling. Likewise, we found that lifestyle features such as diet correlated with taxa that influenced expression of host genes involved in metabolism and cytokine signaling. High-diversity microbiomes altered more host genes than low-diversity ones; when directly compared, low diversity enhanced epithelial barrier-and energy metabolism–related pathways, whereas high diversity activated broader regulatory processes such as RNA processing and receptor-mediated signaling. These findings support the idea that microbiome diversity modulates distinct host physiological states, potentially contributing to inflammatory conditions that are associated with industrialization.

Urban microbiomes have been connected to disease phenotypes in many observational studies [17,20,76]. In general, we observed more interactions with host immunity by urban microbiomes than rural microbiomes, potentially reflecting more immune stimulation in response to urban taxa. Interestingly, these responses were generally attenuated (**Figure 2B, Figure S4**), which may reflect a suppression of immune functions in response to microbial colonization [77]. Another possibility is that these inflammatory responses are modulated by either beneficial or immune-modulating bacteria. In this study, we see evidence for both scenarios: the Bifid shunt, a beneficial carbohydrate fermentation pathway encoded by *Bifidobacterium* [55,58], downregulates TNF signaling in urban microbiomes (**Figure 3G**). *Treponema*, a genus found more abundantly in rural communities (**Figure S2**) [9,78,79], likewise downregulates innate immune functions (**Figure 3C**).

Industrialization has been broadly linked to reduced microbial diversity [17,20,80–82]. Some have hypothesized that this loss of diversity leads to a loss of synchronicity with the host genome [83,84], possibly leading to chronic inflammatory conditions. Here, we observed that high-diversity microbiomes induced more differentially expressed genes than low-diversity microbiomes relative to untreated controls (**Figure 2E, Figure S5**), which may reflect the host encountering a greater variety of microbial cues or metabolic products in more diverse communities. Another theory is that low diversity microbiomes preferentially influence the host through a greater induction of cellular stress compared to high-diversity communities [9,20]. In line with this, we found low-diversity microbiomes to reshape epithelial structure and upregulate energy consumption pathways (**Figure 2F**), a stress response reminiscent of the Warburg effect which characterizes several chronic inflammatory diseases [85,86]. To our knowledge, a direct connection between microbiome diversity and a Warburg-like metabolic shift has not been previously described, although the role of butyrate-producing microorganisms in maintaining epithelial energy balance is well established [87].

Diet and lifestyle are known to shape microbiome composition and are closely linked to health outcomes. Dietary patterns, in particular, are associated with distinct microbial taxa and metabolic functions [9,18,20,40,81] that may influence host phenotype through gene expression. For example, studies have linked increased fiber intake to higher abundances of carbohydrate-degrading microbiota and lower incidence of intestinal disease [18,40,88]. Here, we noticed a similar pattern wherein microbial taxa and functions implicated in carbohydrate fermentation and beneficial short-chain fatty acid production had associations with host diet. This includes mixed acid fermentation, a microbial process with end products including acetate, succinate and lactate [69,89]. We observed that this process inversely correlates with PC1 and influences the expression of pro-inflammatory *IL-15* (**Figure 4F**), which has been implicated in conditions such as obesity, type 1 diabetes and celiac disease [90–93]. Our analyses also supported studies of well-known probiotics: *Bifidobacterium adolescentis*, a short-chain fatty acid-producing species that is decreased in obesity [53,59], correlated with more lactose consumption and was linked to increased expression of *TKT* (**Figure 4B**), a host enzyme which is crucial for connecting glycolysis to downstream metabolic pathways and protecting against intestinal colitis [60].

In fact, some of the organisms that drove expression of the highest number of genes in this analysis, regardless of urbanism status, were well-characterized carbohydrate fermenters and acetate and butyrate producers such as members of *Ruminococcaceae, Lachnospiraceae,* and *Clostridia* (**Figure 3B**). However, these taxa were rarely differentially abundant across populations and showed fewer associations with host lifestyle features (**Figure 4A, Figure S6**). One interpretation is that slight changes in the abundances or activity of more ubiquitous taxa, rather than differentially abundant taxa, may have greater sway over host physiology long-term. This aligns with groups who have postulated that these highly prevalent taxa (*Ruminococcus, Clostridium,* and *Roseburia*) to be among the host’s most ecologically and functionally impactful microbiome constituents [94,95]. Alternatively, individual differentially abundant microbes may exert subtle, targeted effects on host physiology [83,84]. For instance, we recently found that *Bacteroides dorei,* a species enriched in urban microbiomes, may influence immune signaling through modulation of IgA [8] and induction of key inflammatory mediators such as CCN1, despite affecting only a limited number of host genes (**Figure 4C, Figure S2**) [62,63].

While colonocyte systems offer important advantages for studying host–microbiome interactions, such as measurement of epithelium-specific responses, control of environmental factors, and suitability for high-throughput screening, they also have inherent limitations. These include simplified tissue complexity and lack of in vivo physiological conditions. Although primarily cell lines derived from multiple genetic backgrounds can provide broader cellular diversity, this study employed colonocytes from a single genetic background. As such, our findings may not be fully generalizable to populations with diverse genetic profiles. Future experiments could incorporate colonic organoids derived from individuals of diverse ancestries to capture both a broader range of host cell types and explore responses that arise from genetic variation. Finally, our model does not capture long-term, adaptive responses to microbiome variation. Studies conducted in mice have suggested that urban microbiota induces elevated levels of TCRγδ+ T cells [96], demonstrating that microbiome composition can exert lasting immunological effects. To fully elucidate how microbiomes from different communities contribute to metabolic and inflammatory disease, it will be critical to consider both immediate responses in gut epithelium and long-term immune adaptations.

In conclusion, we find that gut microbiomes from urban and rural communities elicit distinct regulatory responses in host colonic epithelial cells, shaped in part by microbial diversity and composition. These findings underscore that while ubiquitous, short-chain fatty acid-producing organisms are highly influential across lifestyles, discrete changes in organisms characteristic of urban communities may influence host immune responses and epithelial cell morphology. Future studies employing organoids or *in vivo* models, particularly those designed to test the effects of individual microbial isolates, will be essential to validate these effects across genetically diverse hosts and to investigate the long-term consequences of microbial composition on immune activation and disease risk in the context of urbanism.

## METHODS

### Sample collection

Stool samples were collected from 58 healthy participants recruited in Ghana, Nigeria, Rwanda and Malaysia as part of the Global Microbiome Conservancy (GMbC) initiative (http://microbiomeconservancy.org). Methods to recruit these participants, collect fecal samples, and sequence microbial metagenomes have been previously described [8,75][75]. Briefly, stool samples were collected in sterile containers and diluted 1:5 in a 25% glycerol solution, then homogenized and immediately flash frozen in 2mL cryogenic tubes. All collections were approved by the relevant local ethics committees: Cape Coast Teaching Hospital Ethical Review

Committee, protocol #CCTHERC/RS/EC/2016/3 and Committee on Human Research, Publication and Ethics of the Komfo Anokye Teaching Hospital, protocol #CHRPE/AP/398/18 (Ghana); Universiti Malaya Medical Research Ethics Committee, MREC ID No.: 2018219-6033 (Malaysia); National Health Research Ethics Committee of Nigeria, protocol #NHREC/01/01/2007-29/04/2018 (Nigeria); National Ethics Committee, protocol IRB 00001497 of IORG0001100 (Rwanda). Urbanism for each community was estimated using the SEDAC Population Estimation Service (https://sedac.ciesin.columbia.edu/mapping/popest/pes-v3/), based on GPS coordinates of the sampling sites. As described previously [8], the total population within a 5 km radius of each locality was quantified as density of inhabitants per square kilometer of land area. Communities with densities >1,000 inhabitants/km² were classified as urban, and those with lower densities as rural. Urbanism was chosen as the variable of interest rather than industrialization in order to avoid sample imbalance issues in downstream analyses.

### Microbiome extraction and storage

In a sterile 0.5% oxygen cabinet, GMbC fecal samples were transferred to a sterile blender cup and combined with 20mL glycerol and 200mL of a 0.90% wt/vol NaCl buffer. Samples were homogenized for 2 minutes, then transferred to a 330-micron filter bag. The liquid suspension filtrate containing the live microbiome was collected, mixed, and aliquoted into vials.

To supplement the sample set of urban samples, 5 microbial suspensions stored in 12.5% glycerol were obtained from healthy USA individuals via OpenBiome (Ind. 1: 02-028-C, Ind. 2: 0065-0016-D, Ind. 3: 0110-0006-01, Ind. 4: 0111-0014-01, Ind. 5: 0112-0002-02). Acquisition and metagenomic shotgun sequencing of these samples has previously been described by our group [30]. GMbC and OpenBiome microbial suspensions were stored at-80°C for downstream use.

### Colonocyte-microbiome co-culture

Colonocyte treatment with live microbiomes has been previously described by Richards et al. [30]. Co-culture was performed using primary human colonic epithelial cells (HCoEpiC, lot 17030), hereby called colonocytes (ScienCell Research Laboratories, Carlsbad, California, USA, 2950). The cells were cultured on plates or flasks coated with poly-l-lysine per manufacturer’s instructions. Cells were cultured in colonic epithelial cell medium supplemented with a growth supplement and Penicillin-Streptomycin per manufacturer’s instructions (ScienCell 2951) at 37°C with 5% CO_2_. 24 hours before treatment, cells were changed to antibiotic-free medium and moved to an incubator at 37°C, 5% CO_2_, and a reduced 5% O_2_.

Live fecal microbiomes were thawed at 37°C the day of the experiment. The microbial density was assessed via a spectrophotometer (Bio-Rad SmartSpec 3000). Colonocytes and microbial cells were combined at a ratio of 1:10 per well in 96 well plates and incubated for two hours at 37 °C, 5% CO_2_, and 5% O_2_. Each well contained a microbiome sample originating from a single individual. Wells containing colonocytes but lacking microbes were cultured in parallel and used as controls.

Following incubation, wells were scraped on ice, pelleted, rinsed with cold phosphate-buffered saline, resuspended in lysis buffer (Dynabeads mRNA Direct kit) and stored at-80°C in preparation for host RNA-sequencing library preparation.

### Host RNA-seq and processing

Extraction of poly-adenylated mRNA from colonocytes was conducted utilizing a Dynabeads mRNA Direct kit (Ambion) per manufacturer’s instructions. As previously described [31], libraries were prepared using a modified NEBNext Ultradirectional library preparation protocol in which barcodes from BIOO Scientific were added by ligation. Libraries were loaded onto two lanes of an Illumina Next-seq 500 in the former Luca/Pique-Regi laboratory at Wayne State University using V2.5 kits for 75 cycles to obtain paired-end reads, 75 bp in length.

Data preprocessing was conducted using the CHURP v. 0.2.2 pipeline from the Minnesota Supercomputing Institute (MSI) [97]. Briefly, reads were analyzed for quality using FastQC v. 0.11.9, trimmed of adapter sequences using Trimmomatic v. 0.40, then analyzed with FastQC once more. Alignment to the human genome was performed using the GRCh38 database and HISAT2 v. 2.2.1. Post alignment, read counts ranged 15,041,596 - 56,944,023, averaged 29,376,099, had a median of 27,624,440 reads, and yielded an average alignment rate of 88.9% across samples (**Figure S1**).

Two outliers, defined by high rRNA contamination and low quality reads, were removed prior to downstream filtering. The R package biomaRt v. 2.54.1 in combination with the Ensembl mart (v.2.22.0) were used to select protein-coding genes and non-novel genes (i.e., no gene symbol). Finally, host genes were selected for downstream analysis if they were expressed (read count > 0) in at least 50% of colonocytes treated with urban microbiomes or 50% of colonocytes treated with rural microbiomes. For subsequent steps (DESeq2, lasso regression, sparse CCA), only the top 3 quartiles of the most variable host genes were considered. This totaled to 14,907 host genes considered for downstream analysis.

### Microbiome DNA-sequencing and processing

Library preparation and microbiome DNA sequencing of GMbC samples has been previously described [75]. Stool samples preserved in RNAlater were subjected to microbial DNA extraction using the MoBio Powersoil 96 kit (now Qiagen Cat No./ID: 12955-4). Genomic DNA libraries were prepared from 1.2 ng of purified DNA using the Nextera XT DNA Library Preparation Kit (Illumina), following the manufacturer’s protocol. Prior to sequencing, libraries were pooled into batches. Insert sizes and concentrations of each pooled library were determined using an Agilent Bioanalyzer DNA 1000 kit (Agilent Technologies). Paired-end sequencing (2×150-bp reads) was performed on a NovaSeq S4 platform (Illumina Inc).

Microbiome metagenomes from GMbC samples and OpenBiome samples were processed together. First, read quality was assessed using FastQC, then trimmed of respective adapters using Cutadapt v. 4.5 [98]. Kneaddata v. 0.12.0 in combination with TRF, Trimmomatic v. 0.39 and Bowtie2 were used to trim repetitive sequences, perform quality trimming, and de-host reads, respectively. Taxonomic profiling was carried out through Kraken 2 [99], utilizing a custom Kraken database that was reconstructed to maximize the mapping rate of metagenomes from rural individuals, and has been previously described [8]. Briefly, the database was constructed using the NCBI taxonomy and genome assemblies retrieved with kraken2-build -- download-library. The names.dmp and nodes.dmp files were manually edited to integrate GMbC isolate genomes (n=6,000; [8,75]) and BIO-ML isolate genomes (n=3,632; [8,100]). These assemblies were added with kraken2-build --add-to-library, and the final reference database was built using kraken2-build --build. Bracken v. 2.7.0 [101] was used for abundance estimation. The abundance table was filtered for bacteria and input into the R package phyloseq v. 1.42.0 for downstream handling [102]. Taxa were retained if they were found at a relative abundance of 0.1% in at least 25% of urban microbiomes or 25% of rural microbiomes. Microbiome metagenomes were agglomerated to separate taxonomic levels, yielding 8, 16, 22, 33, 46 and 135 taxa belonging to the phylum, class, order, family, genus and species classification, respectively. Finally, at each phylogenetic level, samples were subjected to center log ratio transformation.

Functional profiling was performed using HUMAnN3 (v3.8), using the ChocoPhlAn database (v201901_v31) for nucleotide-level mapping, DIAMOND (v2.0.15) for translated protein-level alignment, and UniRef (uniref90_201901b) for annotation. Gene family mapping rate ranged 69.7% to 94.6%, with a median mapping rate of 89.5% and 83.8% for urban and rural microbiomes, respectively. Microbial pathways were first collapsed at the community level (individual microbe contributions were excluded), then filtered to remove unintegrated or unmapped reads. Pathways were retained if they were found at a relative abundance of 0.1% in at least 25% of urban microbiomes or 25% of rural microbiomes, which resulted in 292 pathways. Pathway abundances were subjected to CLR transformation for downstream testing.

### Principal coordinate analysis of microbiome taxonomy

Microbial community composition was analyzed at the genus level using principal coordinates analysis (PCoA). Taxonomic abundances were CLR-transformed, and Euclidean distance was used to compute a sample-wise distance matrix. Ordination was performed using the ordinate() function from the phyloseq package with the “PCoA” method.

### Differential abundance analysis of microbiome taxonomy and function

Differential abundance analysis was conducted using the R package ALDEx2 [103] to evaluate the effects of urbanism on microbial taxa and function. A model matrix was first constructed to include the covariates of microbiome donor age, sex, and country of sampling. For each taxonomic level, we performed centered log-ratio transformation of OTU counts using the function aldex.clr using 128 Monte Carlo samples. Generalized linear model tests were applied using the function aldex.glm to determine associations with urbanism, and effect sizes were estimated with the function aldex.glm.effect. Results were filtered for significance based on FDR < 10%.

### Alpha diversity analysis of microbiome taxonomy

Shannon diversity of all samples was determined using the estimate_richness function from phyloseq [102]. The range of values for Shannon index was 2.20 - 5.42, with a median of 3.62. A student’s t-test (t.test function; base R) was used to test the difference in Shannon diversity values among urban and rural microbiomes, yielding a non-significant *p*-value of 0.369.

In preparation for differential expression analysis, samples with a diversity value less than the median of 3.62 were considered “low diversity”, while the upper half was considered “high diversity” (**Figure 2D**). Both urban and rural samples contributed to the low and high categories; urban microbiomes comprised 41% of the low-diversity subset and 31% of the high-diversity subset (12 urban and 17 rural in “low”, 9 and 20, respectively, in “high”).

### Differential expression analysis of host genes

DESeq2 [104] was used to test for differences in gene expression between untreated control colonocytes relative to those treated with urban or rural microbiomes. As part of the standard DESeq2 pipeline, host read counts were estimated using size factors to control for library depth. Gene-wise dispersions were used to shrink these estimates. To test the effect of urban microbiome treatment versus control and rural microbiome treatment versus control, we employed DESeq2 negative binomial model in conjunction with the Wald test, where colonocyte batch and total RNA reads were included as covariates:

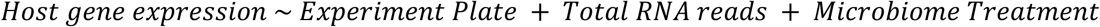

Additional covariates such as microbiome donor age, sex, and country of sampling were ultimately excluded from these models because untreated controls did not receive a microbiome treatment and therefore lacked associated donor metadata, which led to model nonlinearity and convergence issues. Consequently, these variables were only incorporated in subsequent analyses comparing microbiome diversity groups, where controls were excluded.

Contrasts were specified as either “urban microbiome treatment versus control” or “rural microbiome treatment versus control”. *P*-values were adjusted using the Benjamini-Hochberg procedure to control the false discovery rate. Genes with an adjusted p-value below 0.1 were considered significantly differentially expressed. The effect of urban versus rural microbiome treatment was also tested, but was less sensitive and was not subjected to further analysis.

DESeq2 was also utilized to test differences in host gene expression in response to low diversity and high diversity microbiomes. We first employed a similar model:

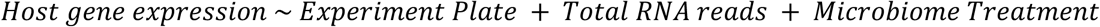

Here, treatment contrasts were specified as either “low diversity microbiome treatment versus control” or “high diversity microbiome treatment versus control”. Genes with an adjusted p-value below 0.1 were considered significantly differentially expressed.

Last, we tested the effect of high-diversity microbiomes versus low-diversity microbiomes on host gene expression. Utilizing the same base model, we incorporated additional microbiome donor covariates of age, sex, and country:

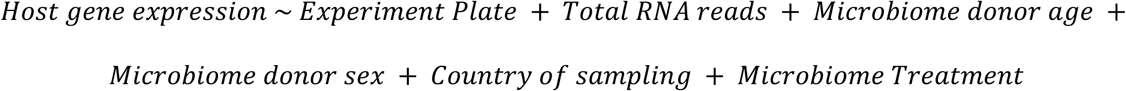

The contrast was specified as “High-diversity microbiome treatment versus low-diversity microbiome treatment”. Genes with an adjusted p-value below 0.1 were considered significantly differentially expressed.

### Over-representation analysis of host gene pathways

To determine the host pathways that were over-represented in significant gene sets, results were first split into upregulated and downregulated genes based on the direction of fold change. The MSigDb databases Hallmark and C2 (including Protein Interaction Database, KEGG, Reactome, and BioCarta) were filtered to retain pathways containing between 25 and 300 genes, to avoid overly narrow or broad categories. Over-representation analysis was performed using the R package clusterProfiler [105], which employs a hypergeometric test to evaluate whether a given pathway is enriched relative to a background set of genes. Pathways with FDR < 10% and at least 3 contributing host genes were considered significantly enriched. The background gene universe was defined as all host genes that passed expression filtering thresholds (see Host RNA-seq and processing, above).

### Lasso regression analysis

Lasso regression analysis was performed in accordance with methods described by Priya et al [24]. Code supporting these steps can be accessed at https://github.com/blekhmanlab/host_gene_microbiome_interactions. Briefly, we applied lasso regression using the R package glmnet (v2.0-13) to model the relationship between microbial abundance and host gene expression, incorporating the covariates Batch, country, age, sex and total reads. We employed leave-one-out cross-validation to estimate the tuning parameter, and inference was performed using the desparsified lasso via the hdi package (v0.1-7), which provided confidence intervals and *p*-values for each predictor. Multiple testing correction was applied using the Benjamini-Hochberg method.

### Sparse CCA analysis

We applied sparse canonical correlation analysis (sparse CCA) to identify multivariate associations between host gene expression and gut microbiome taxonomic profiles, using the PMA package in R (v1.1), following the implementation described by Priya et al (accessed at https://github.com/blekhmanlab/host_gene_microbiome_interactions) [24]. Sparse CCA is a regularized form of canonical correlation analysis that incorporates lasso penalties to enable feature selection in high-dimensional datasets. Prior to model fitting, we performed a grid search over sparsity parameters for both datasets (λ₁ for microbiome features and λ₂ for gene expression) using leave-one-out cross-validation to maximize the correlation between canonical variates. Models were fit using the selected parameters, yielding a series of components, each comprising canonical loading vectors with non-zero weights representing the selected features. For each component, we calculated cross-validated canonical scores and assessed their correlation using cor.test() to evaluate the strength of association. *P*-values were adjusted for multiple testing using the Benjamini-Hochberg procedure to control the false discovery rate.

### Associations of host lifestyle and microbe abundance

Before dimensionality reduction, all lifestyle and dietary questionnaire data were numerically encoded, and features were filtered to retain only those with high inter-individual variance (top 25% by standard deviation) to ensure that downstream analyses focused on the most informative variables. Dietary variables were grouped into interpretable categories based on intake frequency responses. The components of each group were: Processed_Food (CaffeinatedSoda, IndusFruitJuice, IndustrializedCookie, ProcessedCereals, Pasta), Animal_Meat: (AnimalFat, BirdMeat, MammalLivestockMeat), Fish_Meat (FishMeat, Shellfish), Fruits (Banana, PlantainGreenBanana, Avocado, Guava, Oranges, Melon, Mangos, Pomes, Pineapple), Vegetables (Cauliflower, Cucumber, Potatoes, Carrots, SweetPepper, Yams, Okra, Corn, Cabbage, ChiliPepper), Lactose (Milk, Yogurt, Cheese, IceCream), and Legumes (Beans, Lentil). A diet-only principal component analysis was conducted using the prcomp() function in R.

To assess microbiome-lifestyle associations, centered log-ratio (CLR) transformations were performed on taxonomic count data at multiple taxonomic levels and functional count data followed by generalized linear modeling via aldex.glm (ALDEx2), controlling for potential confounders (age, sex, country). This analysis was repeated for all of lifestyle, dietary, blood, and medication features. Features included both aggregate composite scores (e.g., Diet PC1) and raw intake frequencies (e.g., Legumes, Lactose). Significance was determined by evaluating the Holm-adjusted *p*-values derived from the GLM output, with features considered significantly associated if the adjusted p-value was below 0.05.

## Data availability

GMbC shotgun metagenomic, IgA-Seq and human genotyping data are available online on the dbGaP server (Study ID: 38715; Accession: phs002235.v1.p1; Accession: phs002205.v1.p1).

Supporting code can be found at https://github.com/sabrinajarif/gmbc-colonocytes.

## Competing interests

The authors declare no competing interests.

## Author contributions

Conception and design of experiment: S.A, S.N, A.A, F.L, R.B Experiment performance: S.N, A.A, H.M

Data analysis and figure generation: S.A

Writing of manuscript: S.A, with help from S.N, F.L, R.B, M.G, M.P Funding acquisition: M.P, M.G, E.J.A, R.B, F.L

Field administrative work & collection and processing of data and samples: A.A, M.Y.A, S.O.A, Y.A.A, A.D, A.P, Y.N, L.R, K.V, F.I, Y.A.L.L, T.M.P, C.O, J.R, I.E.M, M.P, M.G

All authors read and edited the manuscript.

## Acknowledgments

This work was supported by grants from the Center for Microbiome Informatics and Therapeutics at MIT and the Rasmussen Family Foundation.

This work was supported by NIH grant R35-GM128716 to R.B.

M.P and M.G. received support from the Deutsche Forschungsgemeinschaft (DFG - German Science Foundation) within the Collaborative Research Center (CRC) 1182 on “The Origin and Function of Metaorganisms” (Project-ID 261376515 – SFB 1182, project C5.1 to M.G., project C5.2 to M.P.).

The study received infrastructure support from the DFG (German Science Foundation) within the Cluster of Excellence 2167 “PrecisionMedicine in Chronic Inflammation (PMI)” (EXC 2167-390884018).

M.G. received funding from the European Research Council (ERC) under the European Union’s Horizon 2020 research and innovation programme (CoG VESICULOME, Grant agreement No. 101126254).

M.G. and M.P. received funding from the DFG – Project number 426660215 with the RU 5042 miTarget on “The microbiome as a therapeutic target in inflammatory bowel disease”, subprojects TP01 “Targeting intestinal yeasts and pathogenic yeast-responsive CD4+ T cells in Crohn’s disease” and TP02 “Ecology and function of synthetic bacterial communities for the understanding and modulation of IBD-associated microbiomes”.

The project received funding from the European Union under the Horizon Europe grant agreement No. 101095470 (project miGut-Health). Views and opinions expressed are however those of the author(s) only and do not necessarily reflect those of the European Union nor European Health and Digital Executive Agency (HaDEA). Neither the European Union nor HaDEA can be held responsible for them.

## SUPPLEMENTARY FIGURES

**FIGURE S1.**
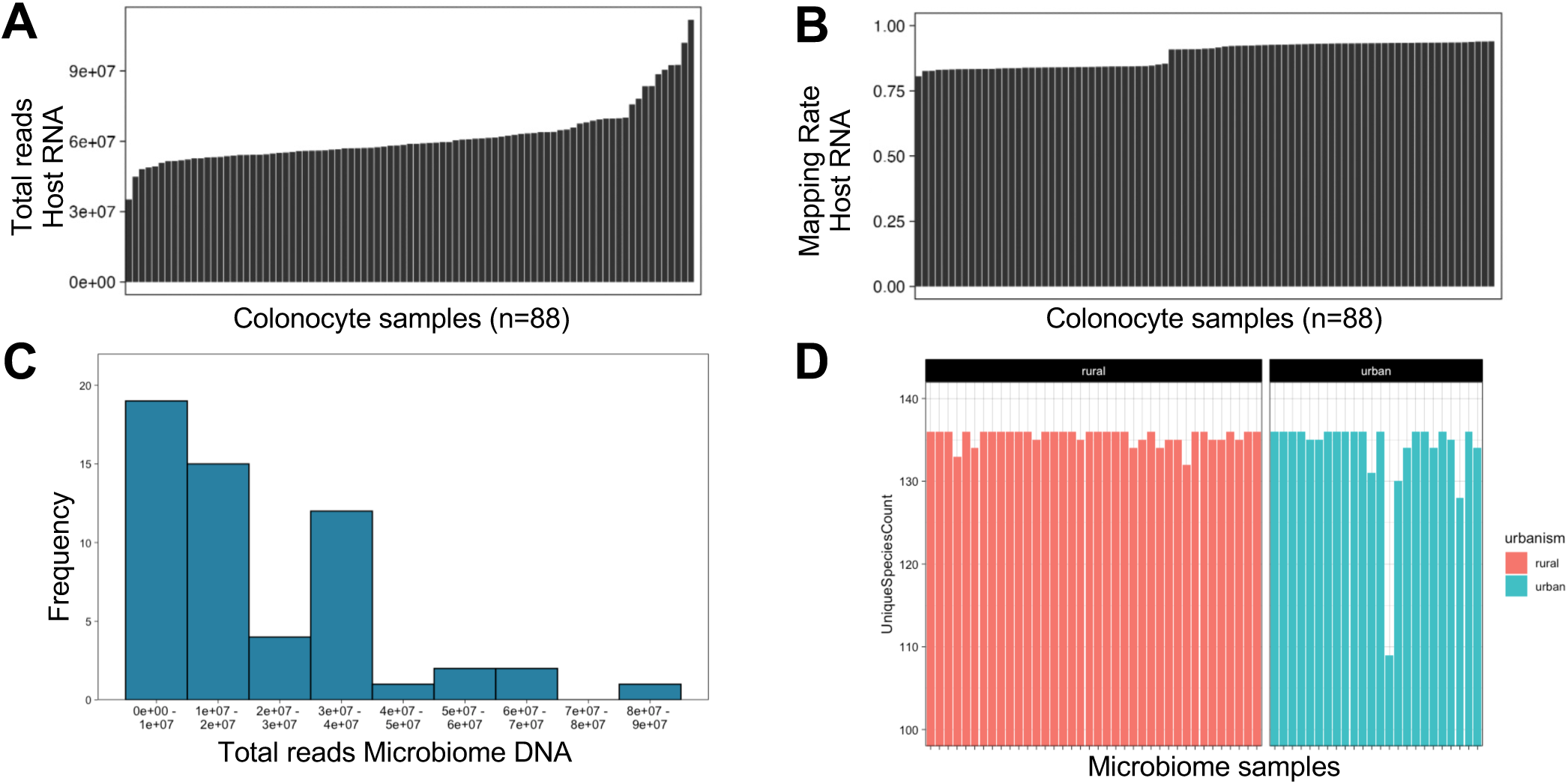
Relevant for. **Figure 1 and Methods. QC and differential abundance of microbiome data. A** Total reads for host colonocyte RNA-sequencing **B** Mapping rate of host colonocyte RNA-sequencing **C** Total reads for metagenomic shotgun sequencing, ranging 3.5 to 87 million reads. **D** Total number of unique microbial species assigned to each sample.

**FIGURE S2.**
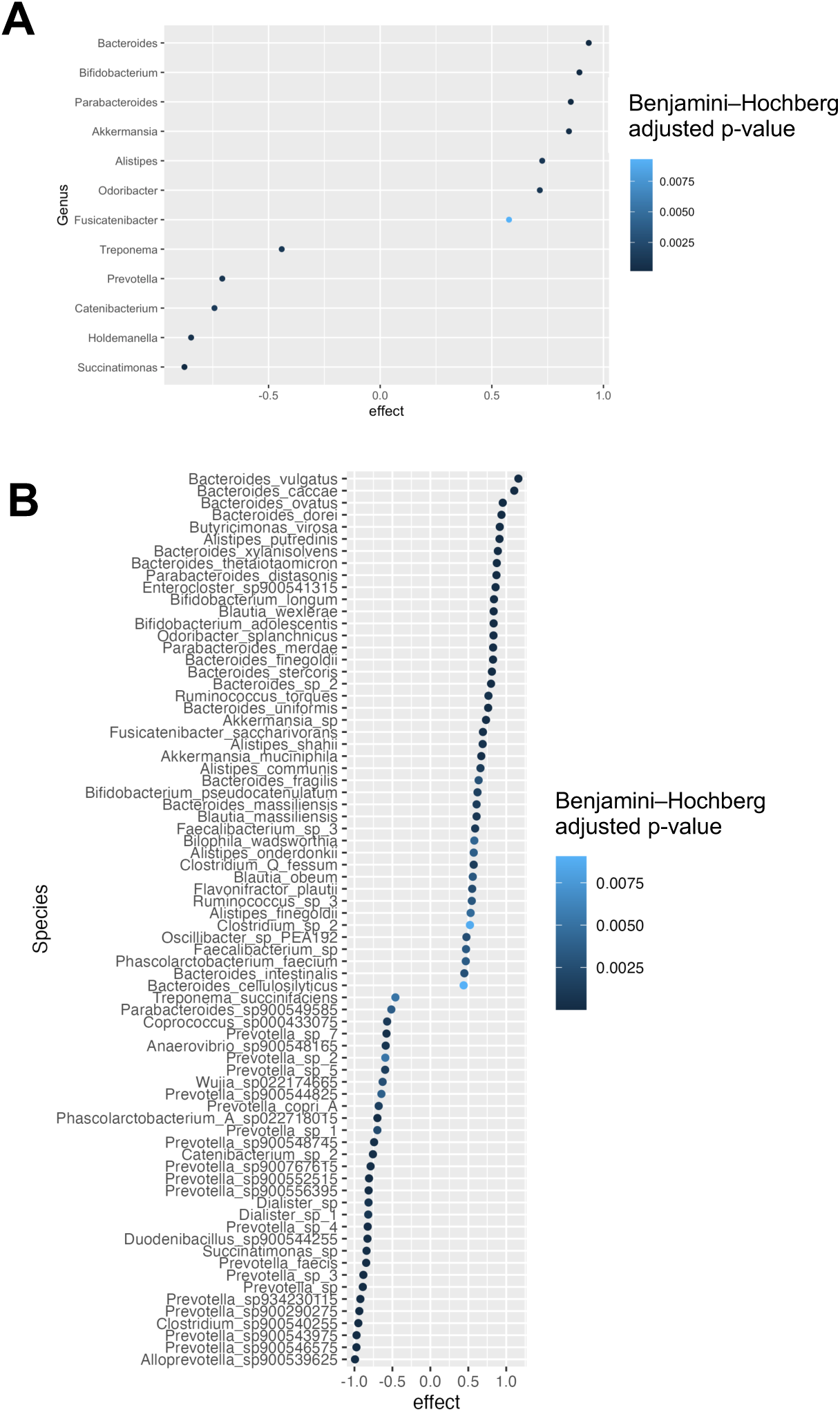
Relevant for. **Figure 1. Taxonomic differential abundance. A**, **B** Differential abundance analysis of microbiome samples in urban v. rural populations, shown at the genus (A) and species (B) level. Effects > 0 reflect taxa that are enriched in urban populations, < 0 are those enriched in rural populations. Only significant (FDR < 0.1) results are shown.

**FIGURE S3.**
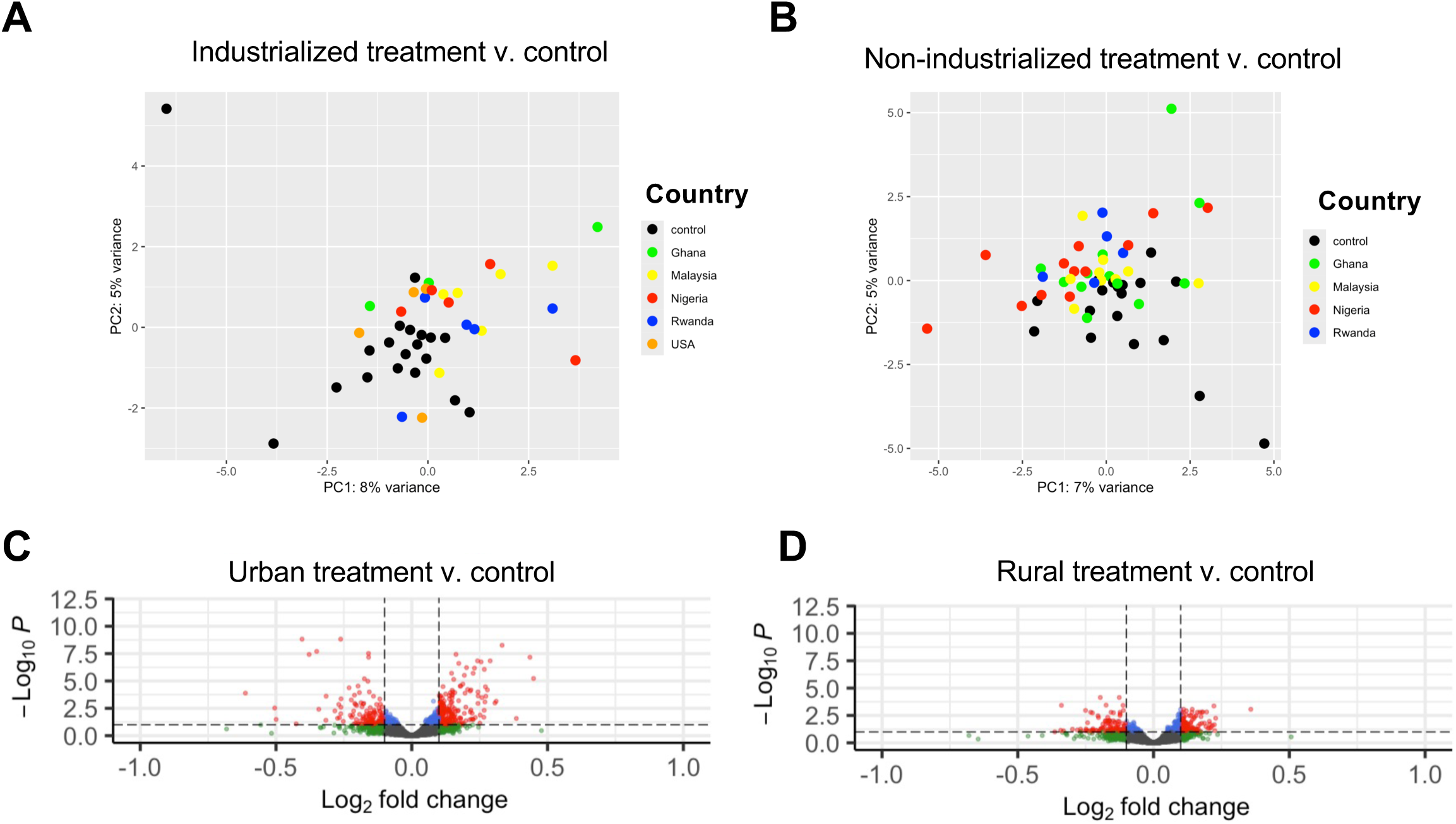
Relevant for. **Figure 2. A, B** PCA plot depicting gene expression of colonocyte samples. Black dots indicate untreated controls. A, colored dots represent colonocytes treated with urban microbiomes from Ghana, Malaysia, Nigeria, Rwanda, and the USA (green, yellow, red, blue, orange, respectively). B, same as A, but colored dots are colonocytes treated with rural microbiomes. **C, D** Volcano plots showing host gene expression in response to urban microbiome versus control (C) and rural microbiome treatment versus control (D). Log2 fold change value (x axis) and *p-*value (y axis) results from differential expression analysis between treatments.

**FIGURE S4.**
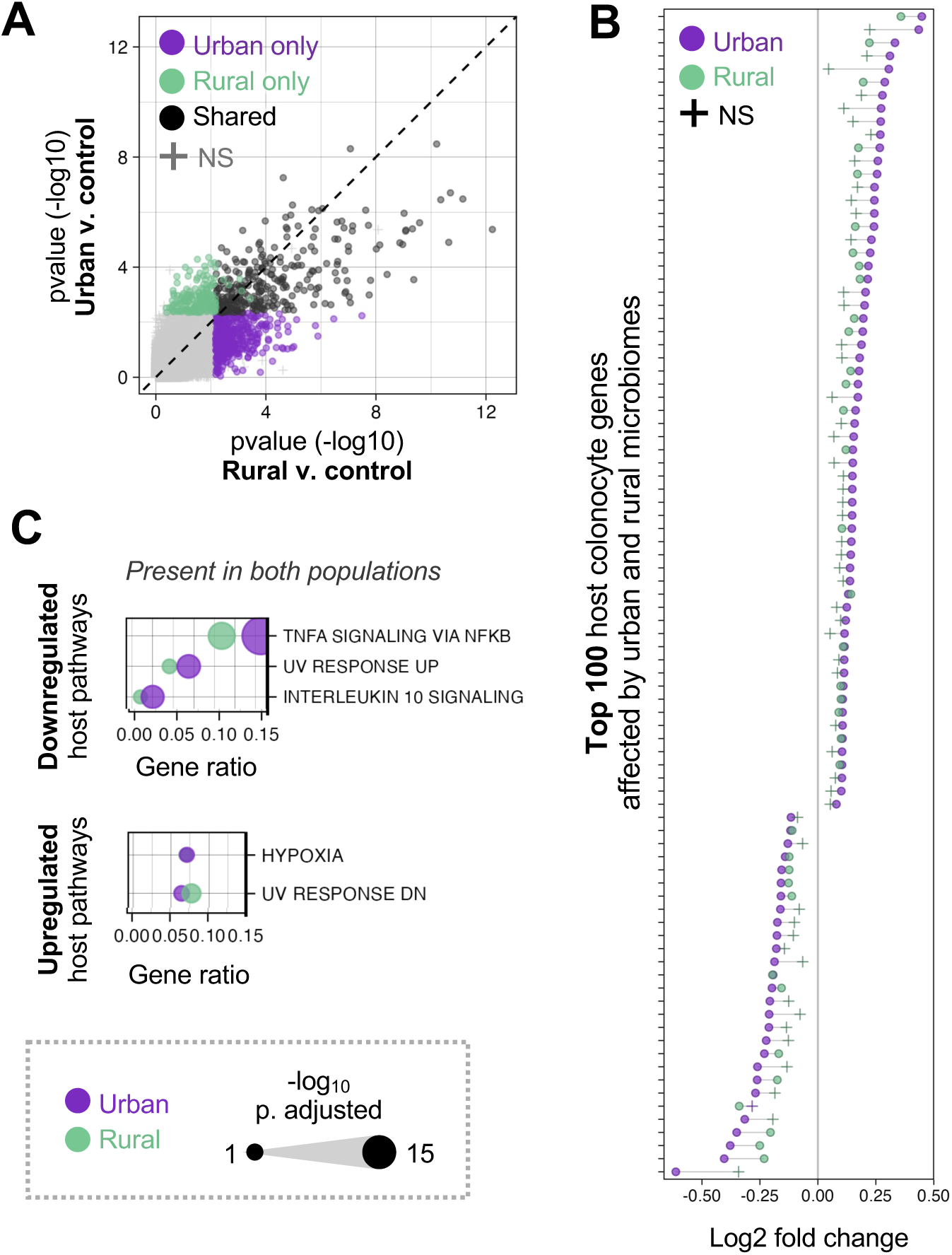
Relevant for Figure 2. **A DEGs compared to control condition.** Left: Raw p-values of host DEGs in response to urban (x-axis) and rural (y-axis) microbiomes. Dots indicate significant DEGs (FDR < 1×10-1) in urban only (purple), rural only (green), or both urban and rural (black). **B Top 100 host DEGs**. Subset of 100 host genes with lowest FDR-corrected *p* values and greatest absolute log2 fold changes in response to either urban (purple) or rural (green) microbiomes. Circles indicate significant (FDR < 1×10^-1^) genes, crosses represent not significant (NS) genes. **C Significant (FDR < 1×10^-1^, gene count ≥ 3) host gene pathways enriched in response to microbiome conditions.** Pathways that are found enriched in response to <u>both</u> urban and rural treatments shown. Down-and upregulated pathways are shown in the top and bottom panels, respectively. Dot size indicates-log10 (*p*-value). Urban and rural treatments produced equal values for the gene ratio for the host pathway of hypoxia.

**FIGURE S5.**
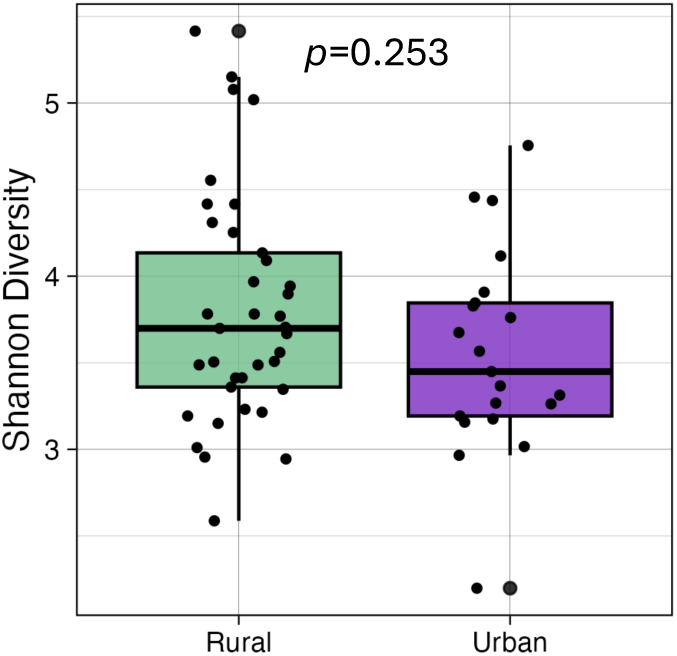
Relevant for Figure 2. Shannon diversity values among rural and urban microbiomes, respectively. Shannon diversity did not differ significantly between rural and urban microbiomes (Wilcoxon rank-sum test, *p* = 0.253).

**FIGURE S6.**
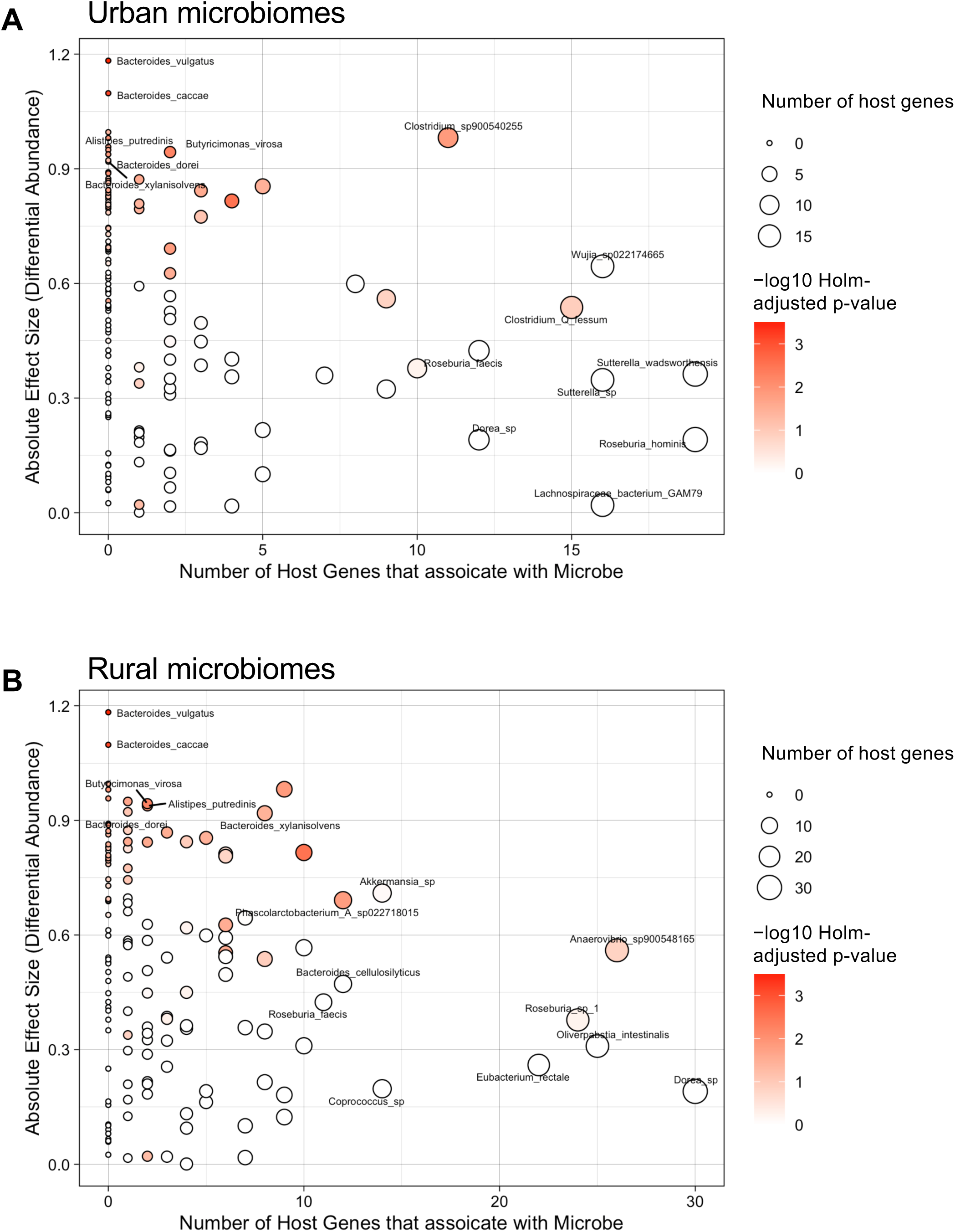
Relevant for Figure 3. Relationship between differential abundance of microbes in urban and rural microbiomes and their effect on host gene expression. **A** Circles represent all microbes found in urban microbiomes. x axis: absolute effect size from differential abundance calculation. y axis: number of host genes that each microbe associates with in pairwise lasso analysis. Shading of circle corresponds to holm adjusted *p*-value of differential abundance analysis. Microbes found in lower right quadrants have strong effects on host gene expression, but are not differentially abundant among urban and rural populations. Microbes found in top left quadrant are differentially abundant, but have minimal impact on host gene expression. **B** Same as A, but depict all microbes found in rural microbiomes.

**FIGURE S7.**
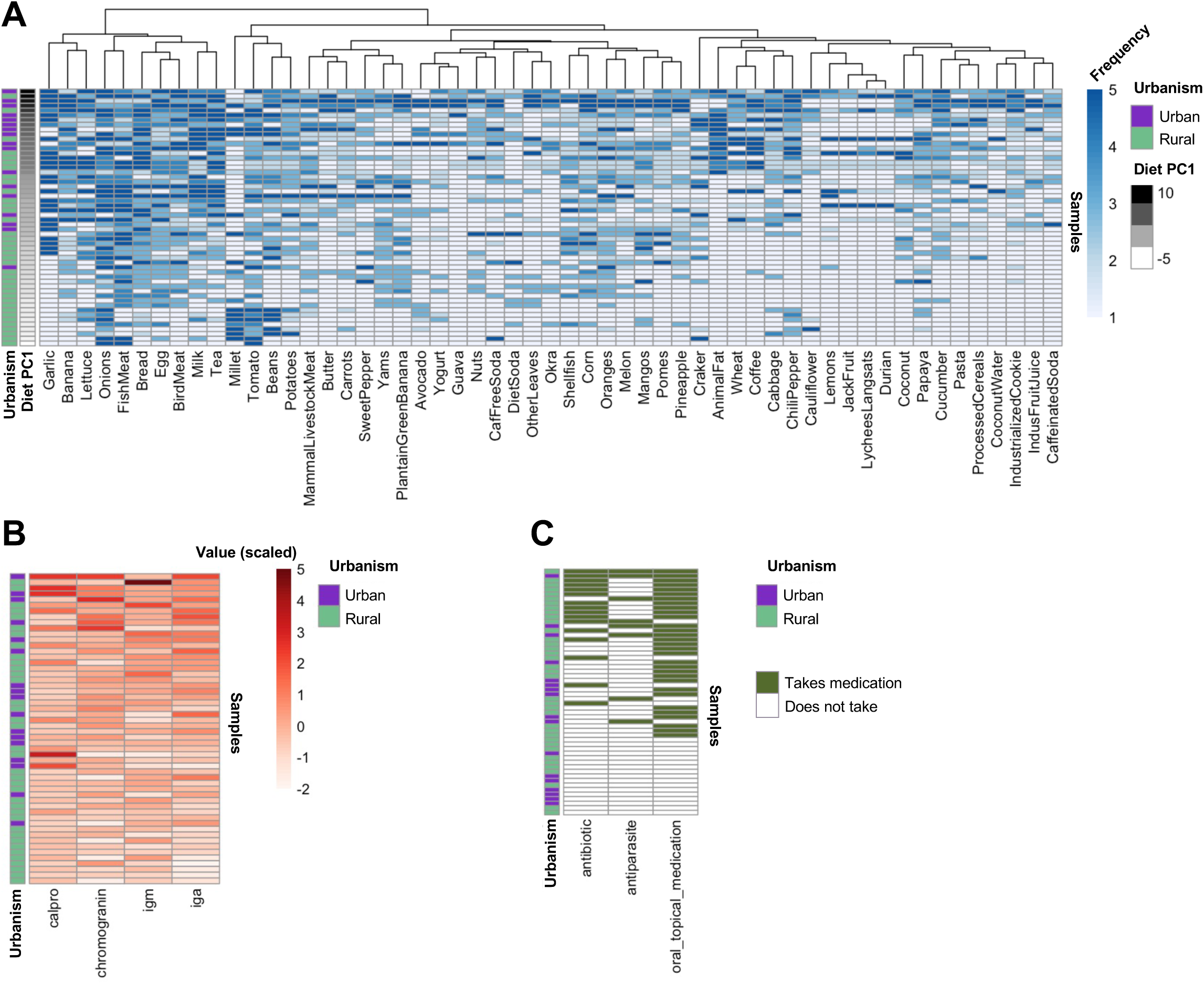
Relevant for Figure 4. **A** Heatmap of diet questionnaire responses (x axis) per participant (y axis). Frequency of consumption indicated by shading of blue tile. Hierarchical clustering applied to x axis. Further annotations provided in left panels: diet diversity (number of types of food consumed at least once), urbanism (green=rural, purple=urban), and value of PC1 diet. **B** Heatmap of blood markers (x axis) per participant (y axis). Scaled value of blood marker concentration indicated by shading of red tile. Further annotations provided in left panels: zscore of summed scaled blood marker value, urbanism. **C** Heatmap of medication use (x axis) per participant (y axis). Participants who take medication indicated by green tile; those who do not take medication indicated by white tile. Further annotations provided in left panels: medication diversity (number of classes of medications used by participant) and urbanism.

**FIGURE S8.**
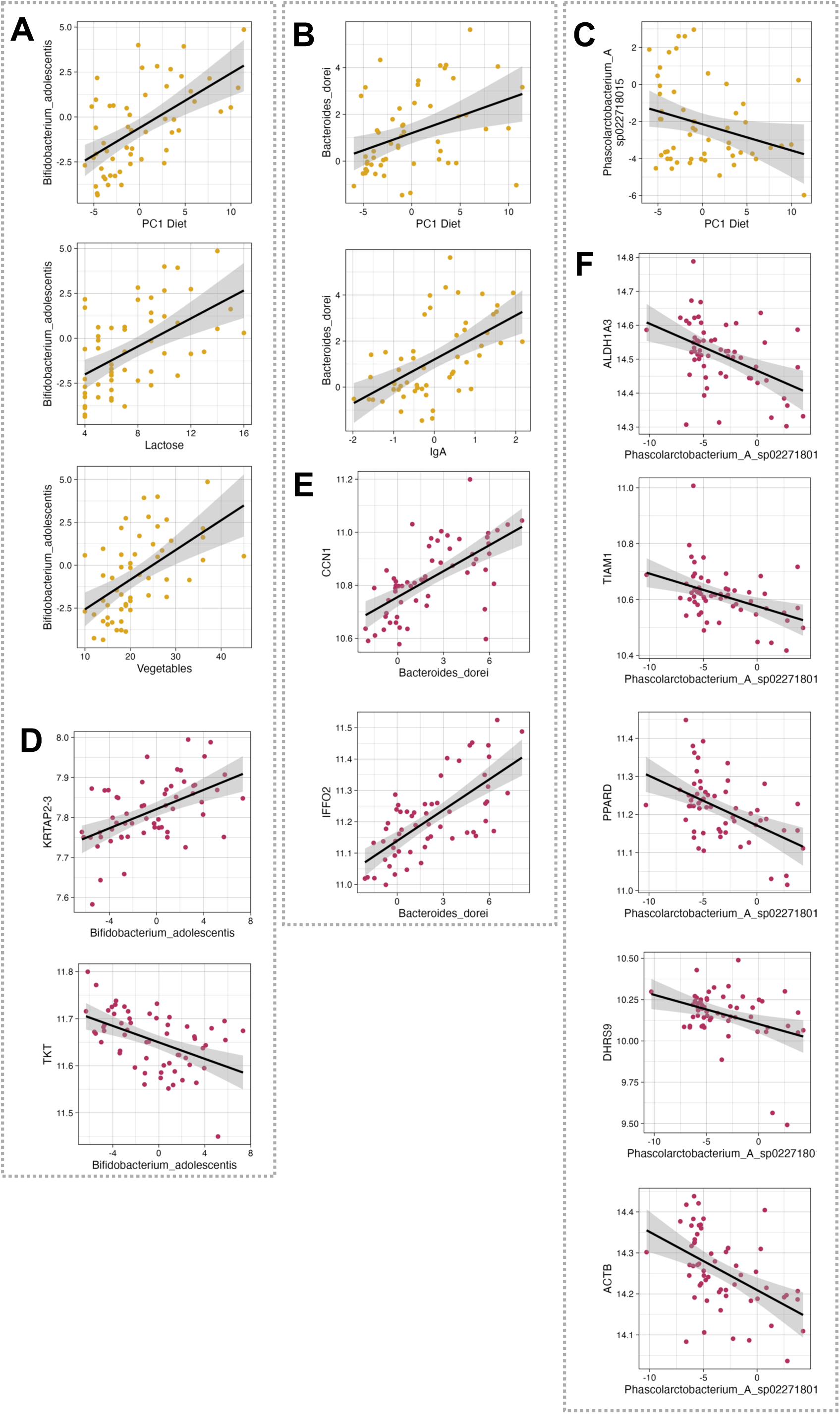
Relevant for Figure 4. **A-C** Scatterplot of lifestyle frequency (x-axis; **Methods**) and microbe abundance (CLR-transformed; y-axis) for *Bifidobacterium adolescentis, Bacteroides dorei,* and *Phascolarctobacterium A sp022718015,* respectively. **D-F** Scatterplot of microbe abundance (CLR-transformed; x-axis) and host gene expression (DESeq2 VST-transformed expression, **Methods**; y-axis) for *Bifidobacterium adolescentis, Bacteroides dorei,* and *Phascolarctobacterium A sp022718015,* respectively.

## SUPPLEMENTARY TABLES

**TABLE S1.** Relevant for Figure 1. Participant metadata.

| database | country | ethnicity | n urban | n rural | n male<br>n female | age<br>range |
| --- | --- | --- | --- | --- | --- | --- |
| gmbc | Malaysia | jahai;thai;batek;chinese<br>;indian | 6 | 9 | 7_8 | 18_59 |
| gmbc | Rwanda | rwandan | 5 | 5 | 6_4 | 24_54 |
| gmbc | Ghana | fante;ashanti;ga;ga_na;<br>akan_fante;akan_akua<br>pem | 3 | 12 | 5_10 | 23_55 |
| gmbc | Nigeria | yoruba;igbo;ibibio;efik | 5 | 12 | 9_8 | 19_59 |
| openbio<br>me | USA | unknown | 5 | 0 | unknown | unknow<br>n |

**TABLE S2.**
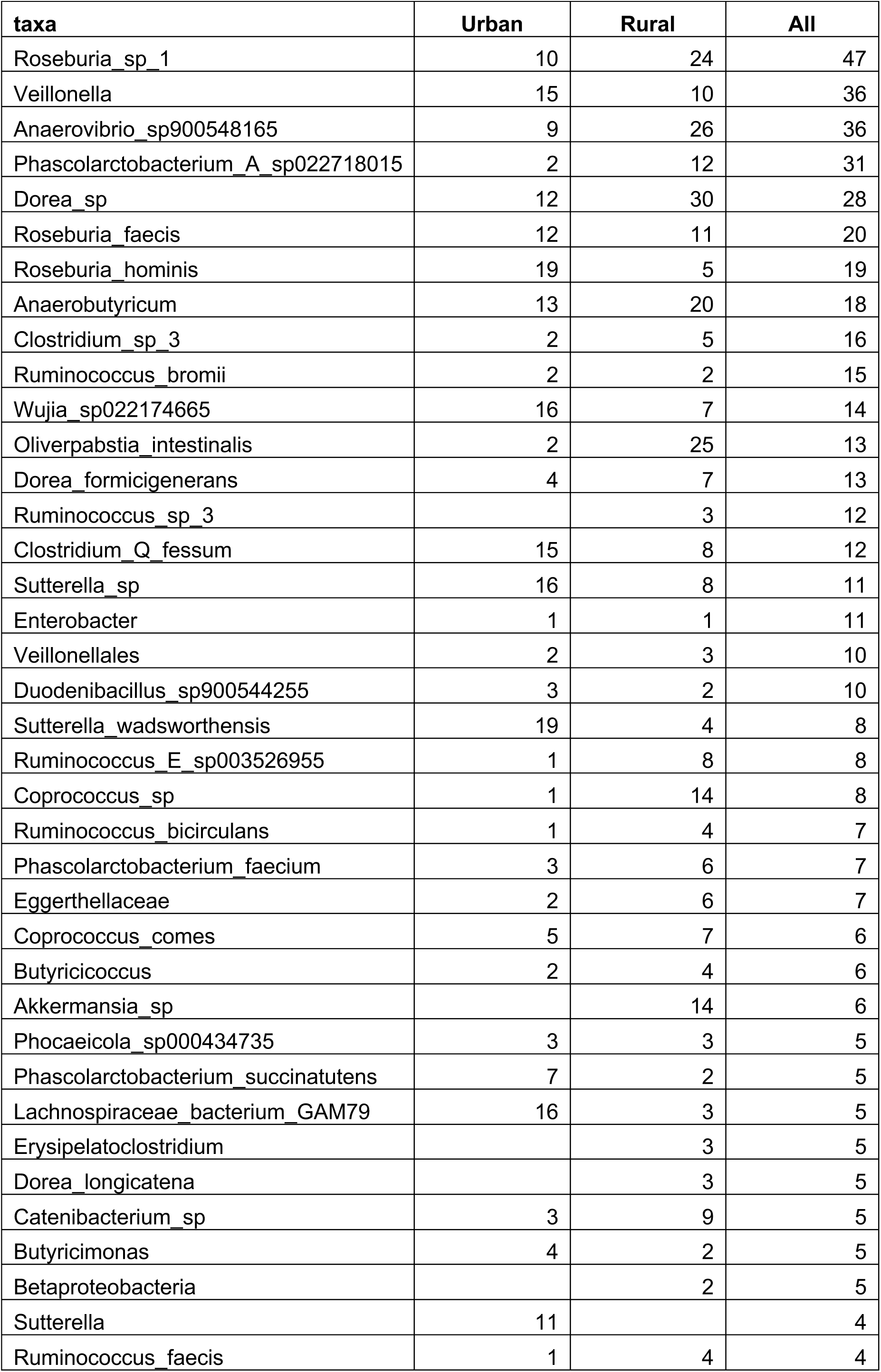
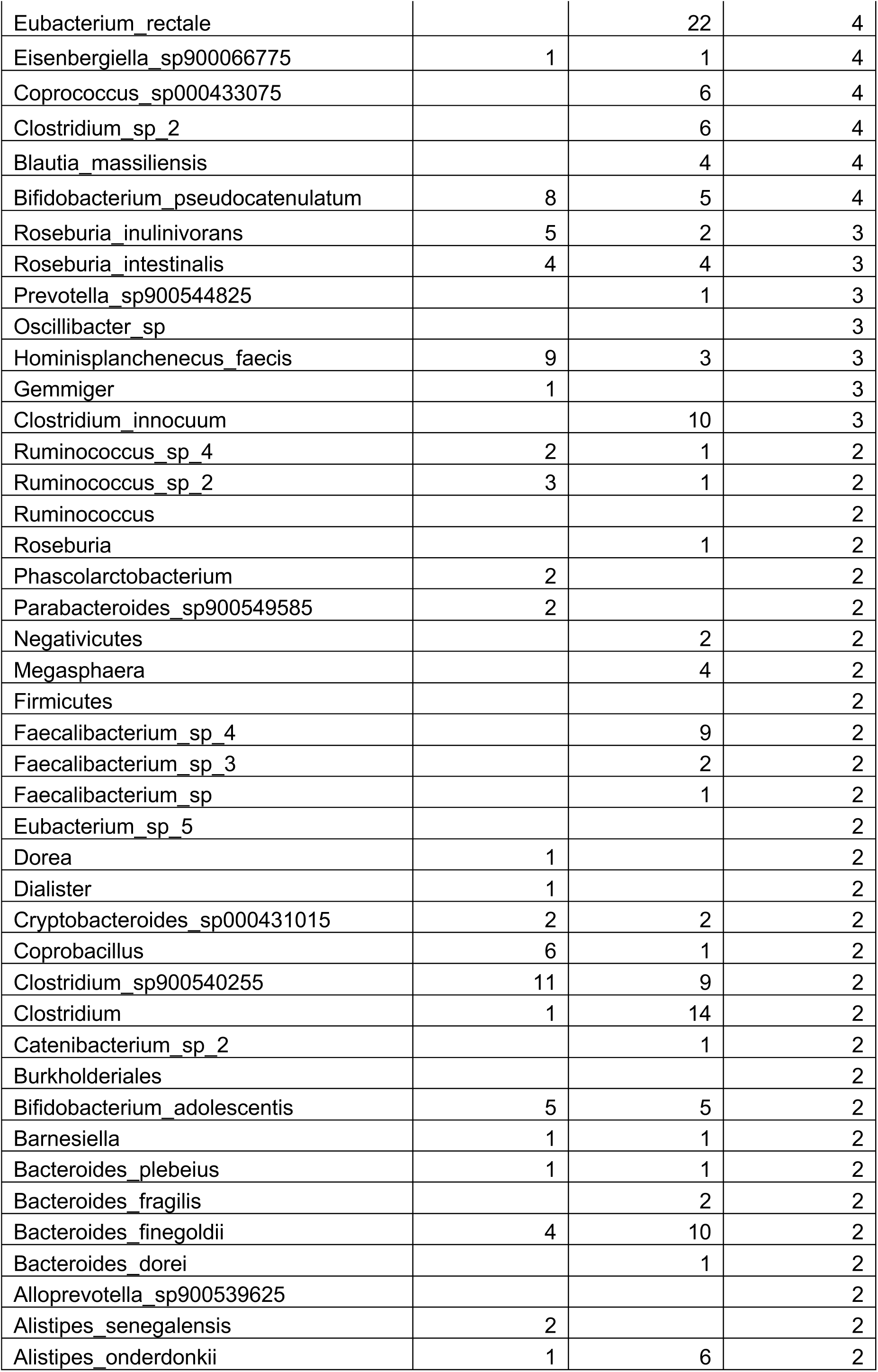
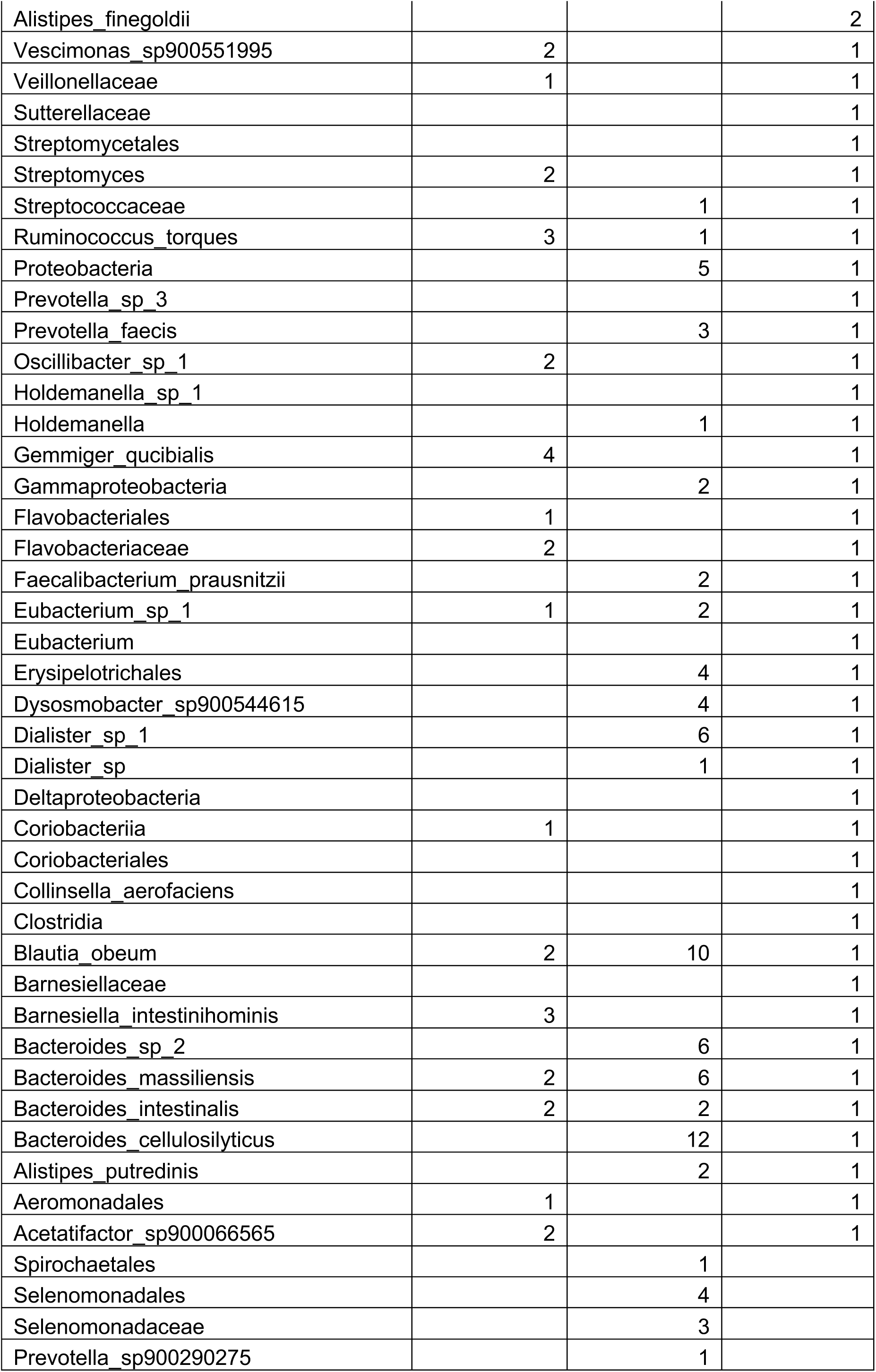
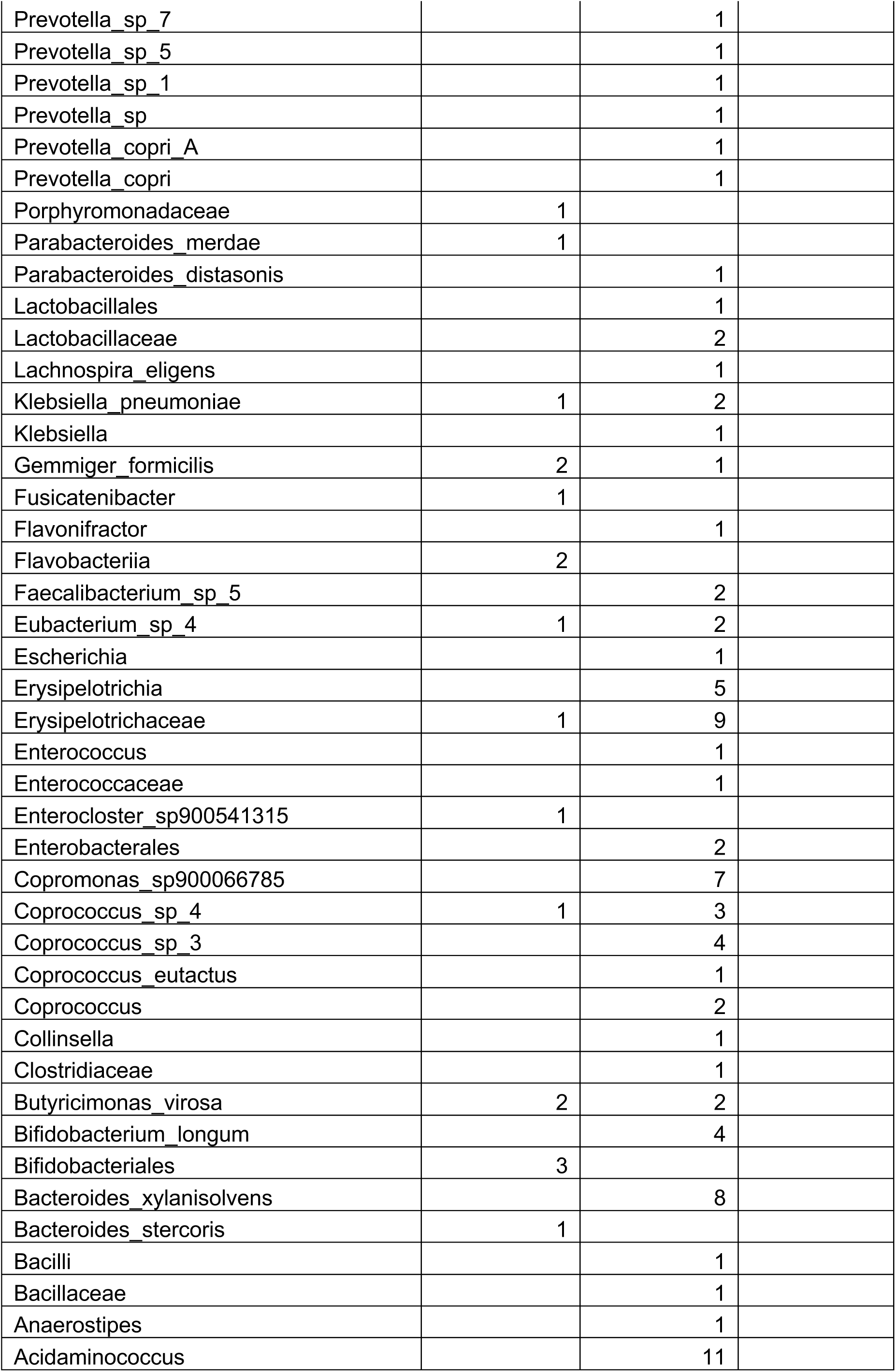
Relevant for Figure 3. Number of associations of microbes with host genes.

**TABLE S3.**
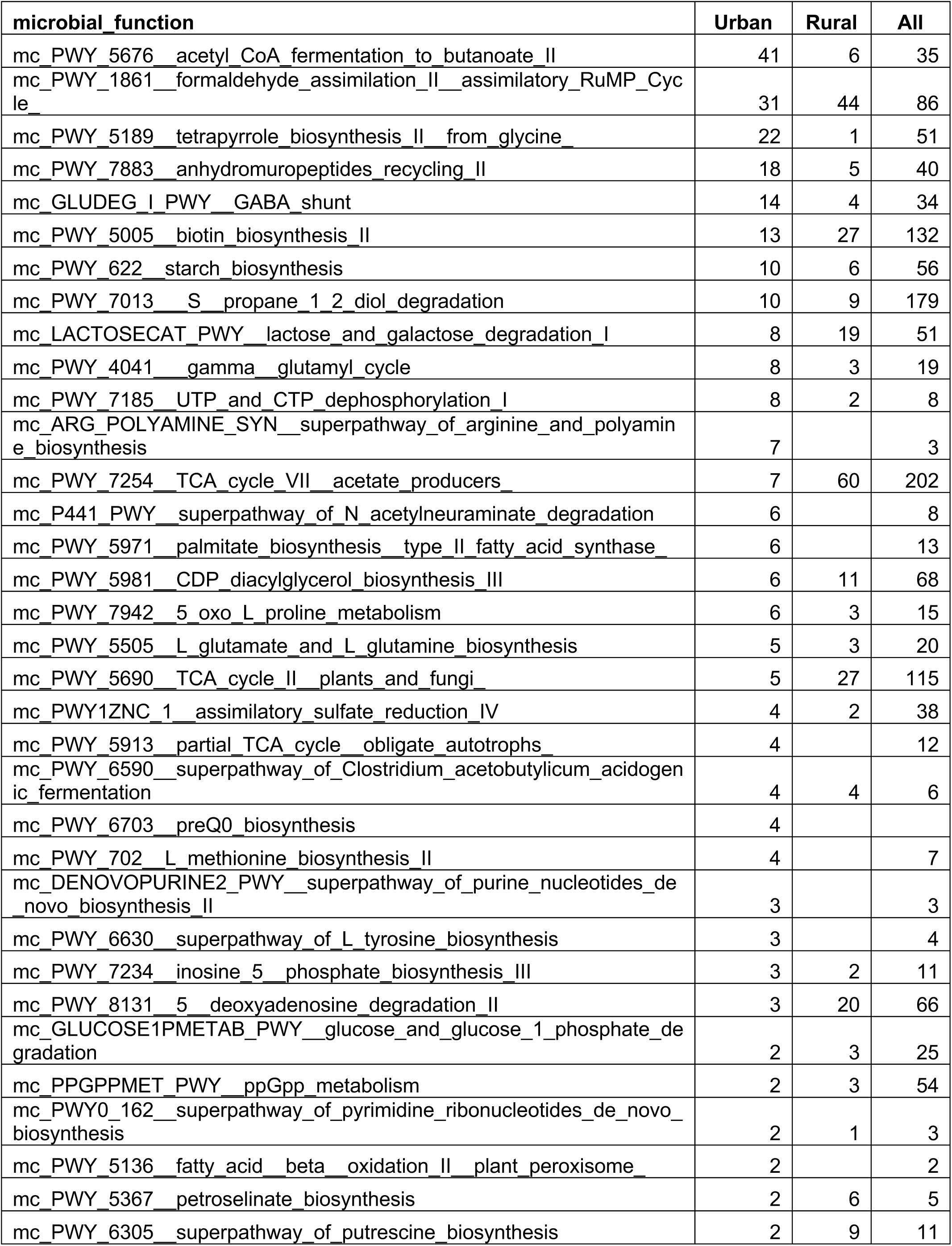
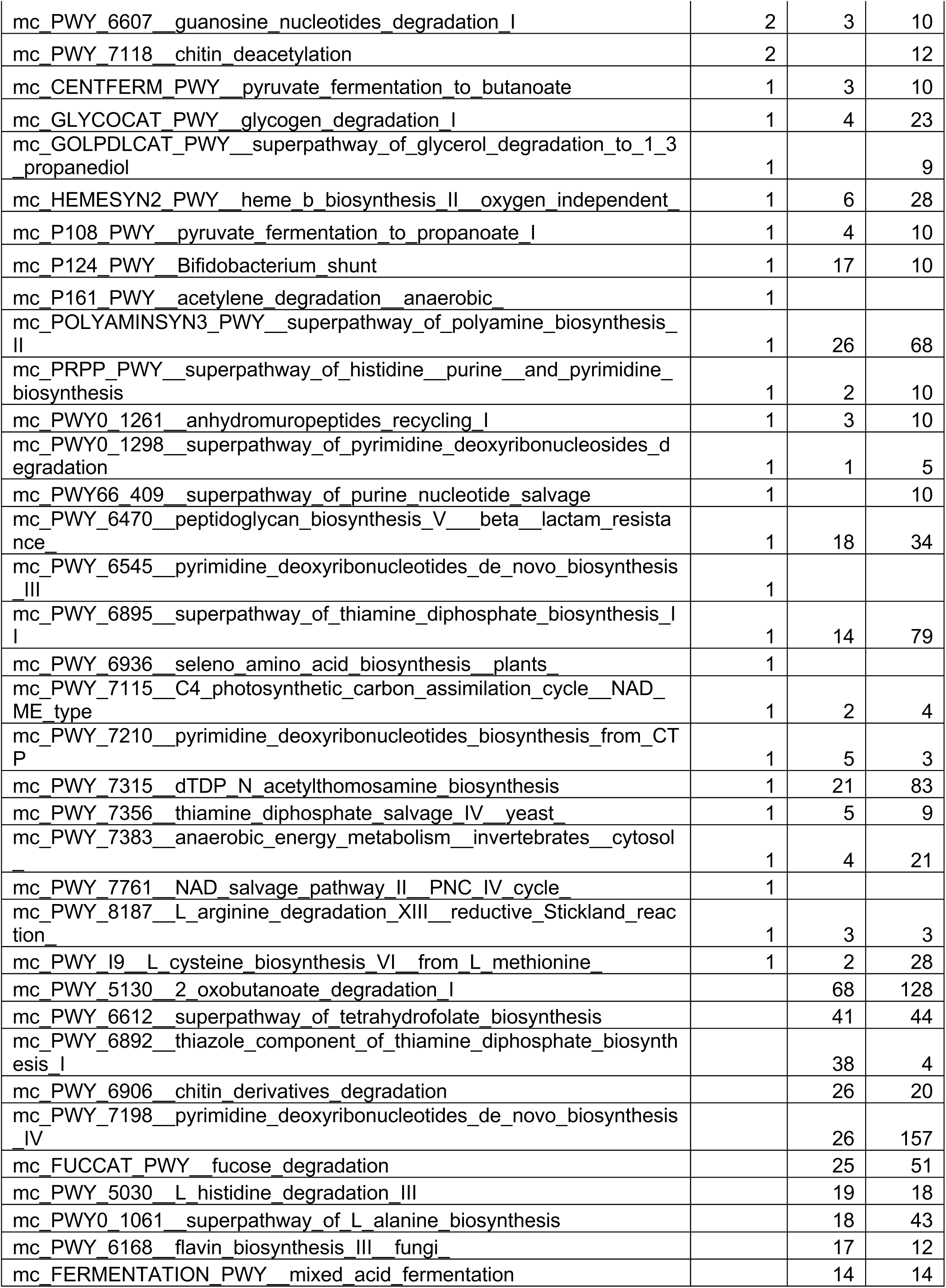
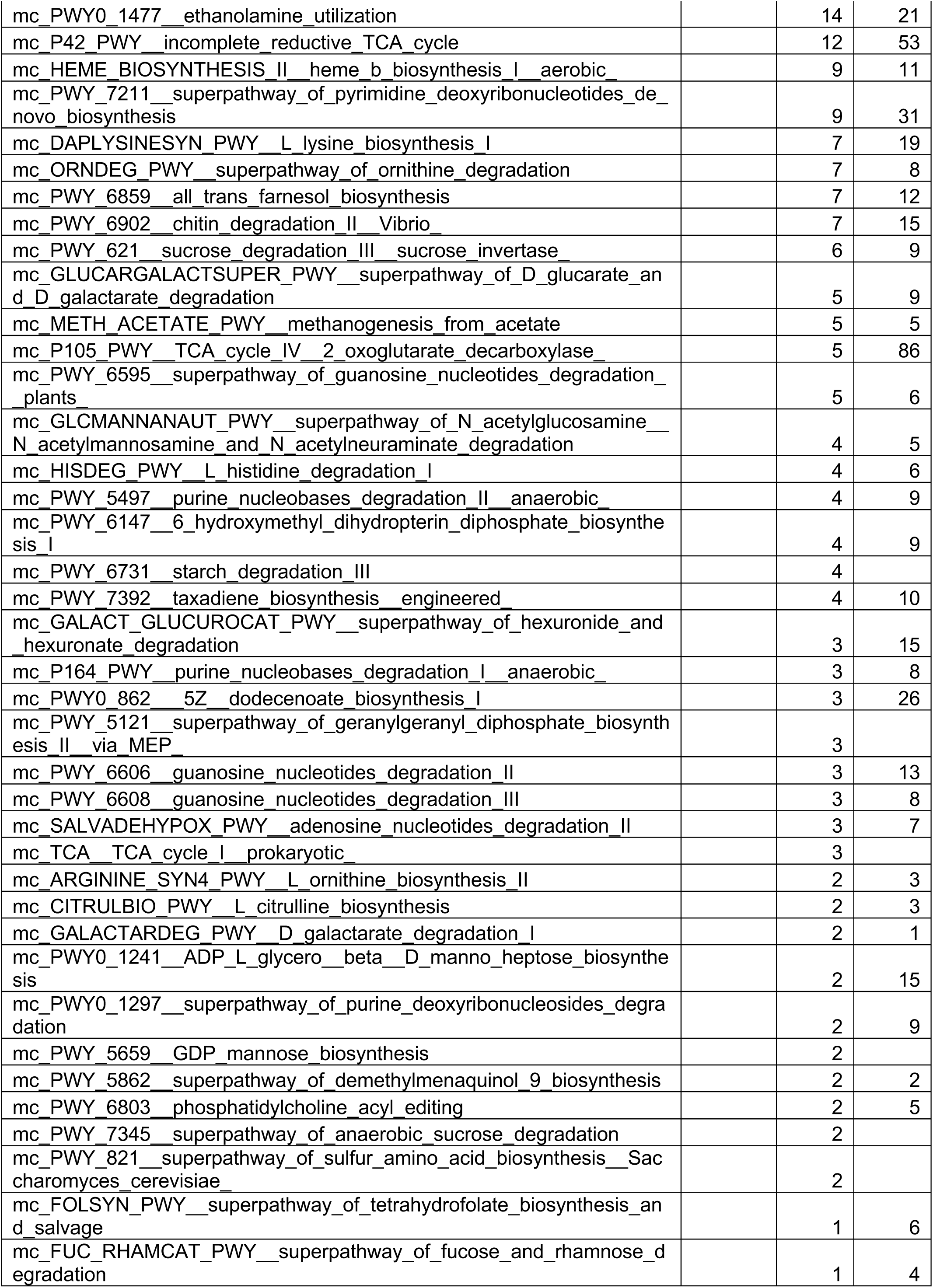
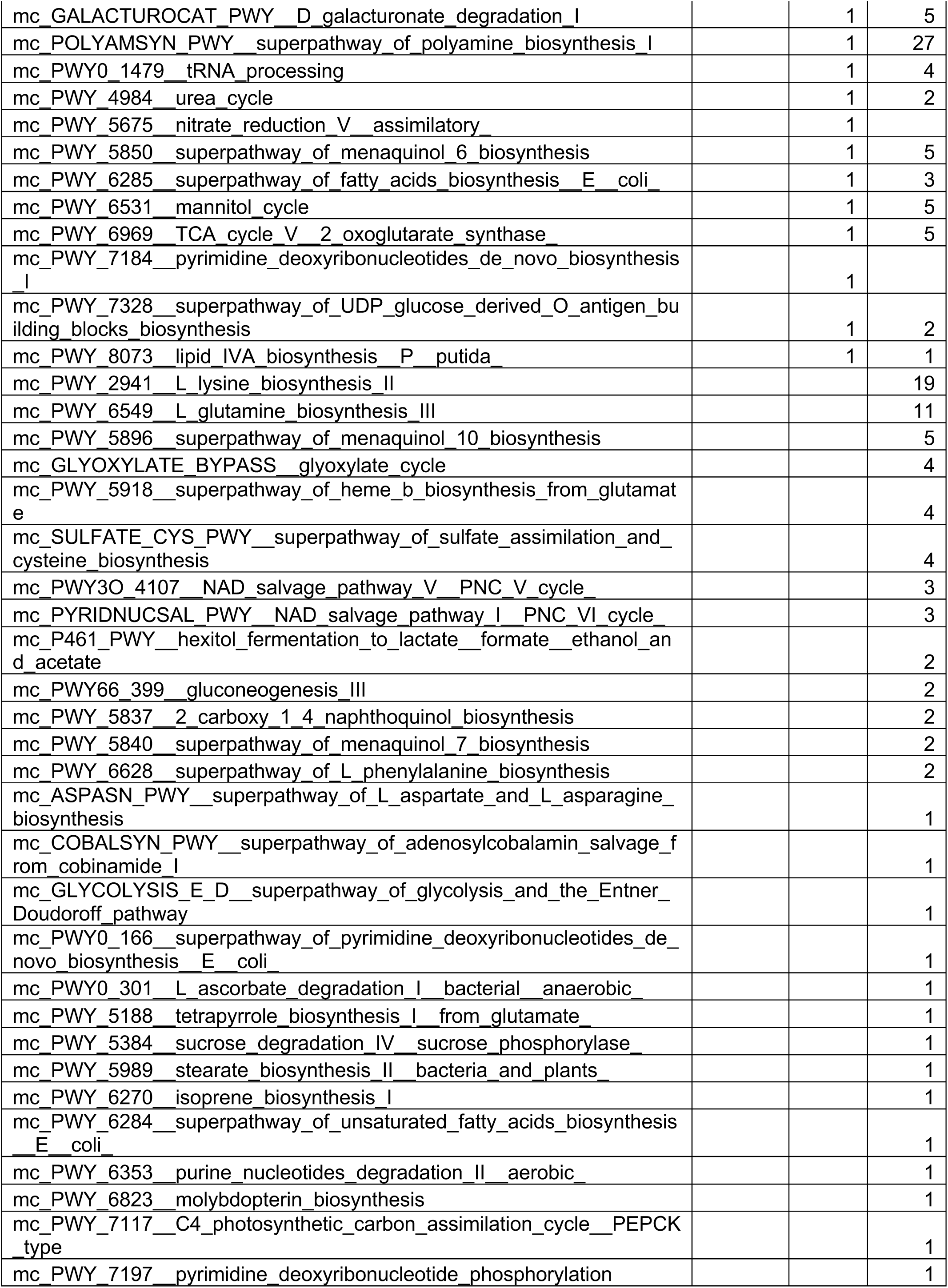
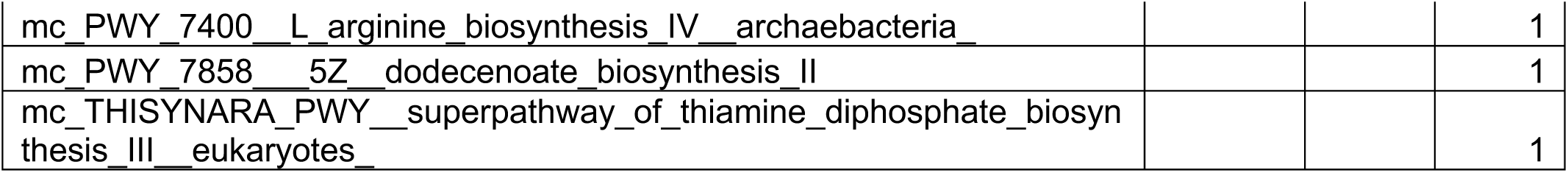
Relevant for Figure 3. Number of associations of microbes with host pathways.

**TABLE S4.** Relevant for Figure 3. Clusters of correlated host genes/pathways and microbes.

| **Host gene & Microbes** |  |  |
| --- | --- | --- |
| **Urban** |  |  |
| *Cluster* | *microbe_taxa_cluster* | *n_host_genes* |
| 2 | Spirochaetaceae; Treponema | 377 |
| 3 | Coprobacillus; Roseburia_sp_1 | 386 |
| 4 | Flavobacteriia; Flavobacteriaceae | 370 |
| 5 | Klebsiella; Klebsiella_pneumoniae | 390 |
| **Rural** |  |  |
| *Cluster* | *microbe_taxa_cluster* | *n_host_genes* |
| 1 | Akkermansiaceae; Akkermansia | 389 |
| 2 | Bacteroides_finegoldii; Parabacteroides_merdae | 399 |
| 3 | Clostridiaceae; Clostridium | 397 |
| 4 | Erysipelotrichia; Erysipelotrichales | 384 |
| 8 | Enterobacter; Klebsiella; Anaerobutyricum;<br>Klebsiella_pneumoniae | 386 |
| 9 | Dialister; Dialister_sp | 394 |
| 10 | Coprococcus; Coprococcus_sp | 405 |

| **Host gene & Microbe pathways** |  |  |
| --- | --- | --- |
| **Urban** |  |  |
| *cluster* | *microbe_function_cluster* | *n_host_genes* |
| 3 | P124_PWY_Bifidobacterium_shunt;<br>mc_PWY_5676_acetyl_CoA_fermentation_to_butanoate_II | 221 |
| 4 | PWY_7118_chitin_deacetylation;<br>PWY_8131_5_deoxyadenosine_degradation_II | 231 |
| 10 | PPGPPMET_PWY_ppGpp_metabolism | 239 |
| **Rural** |  |  |
| *cluster* | *microbe_function_cluster* | *n_host_genes* |
| 1 | HSERMETANA_PWY_L_methionine_biosynthesis_III;<br>PWY_7221_guanosine_ribonucleotides_de_novo_biosynthesis;<br>PWY_7977_L_methionine_biosynthesis_IV | 233 |
| 4 | PPGPPMET_PWY_ppGpp_metabolism;<br>PWY_6731_starch_degradation_III | 231 |
| 6 | ARGININE_SYN4_PWY_L_ornithine_biosynthesis_II;<br>GLUDEG_I_PWY_GABA_shunt | 247 |
| 9 | P164_PWY_purine_nucleobases_degradation_I_anaerobic_;<br>PWY0_862_5Z_dodecenoate_biosynthesis_I | 231 |
| 10 | PWY_6607_guanosine_nucleotides_degradation_I | 236 |

**TABLE S5.**
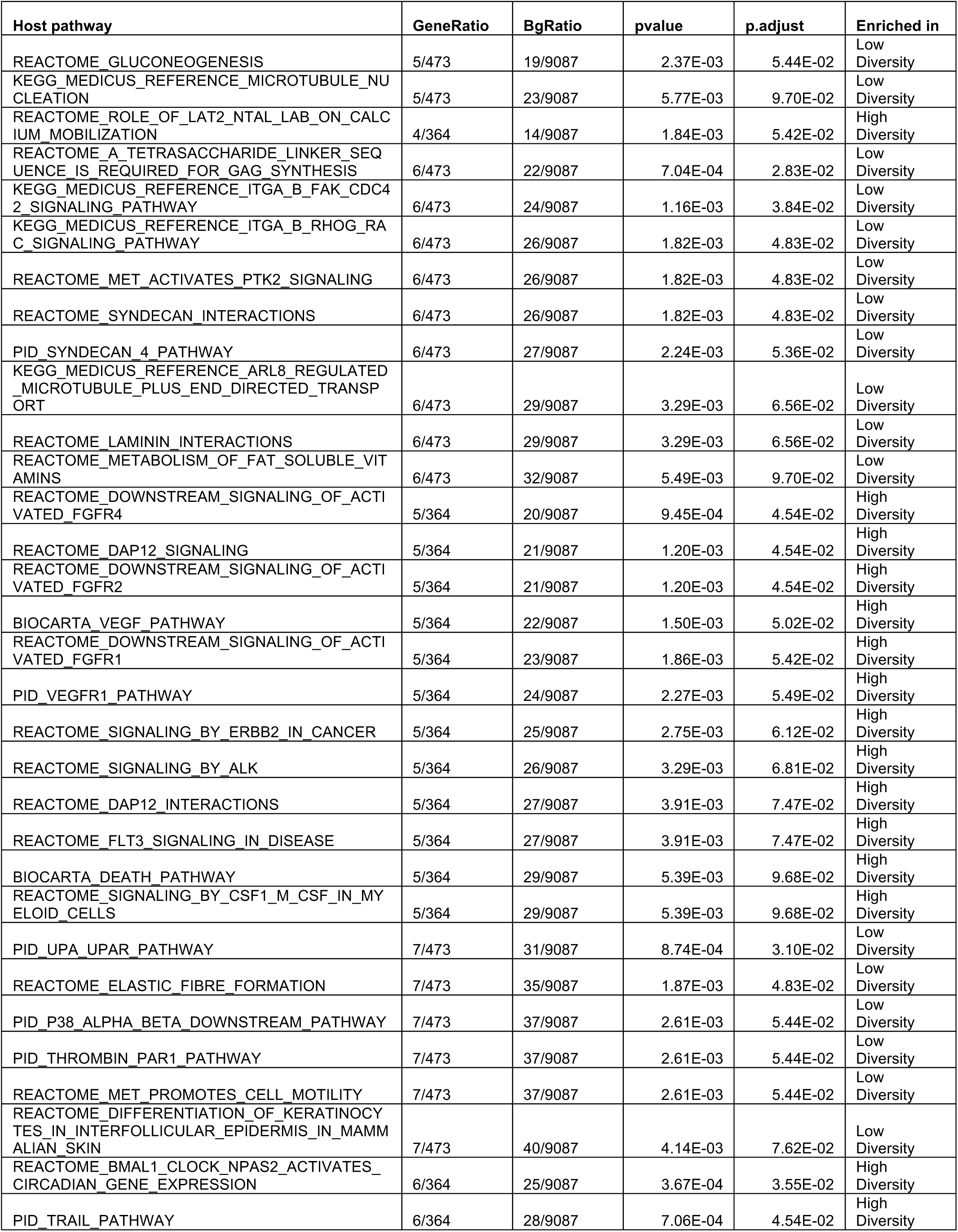
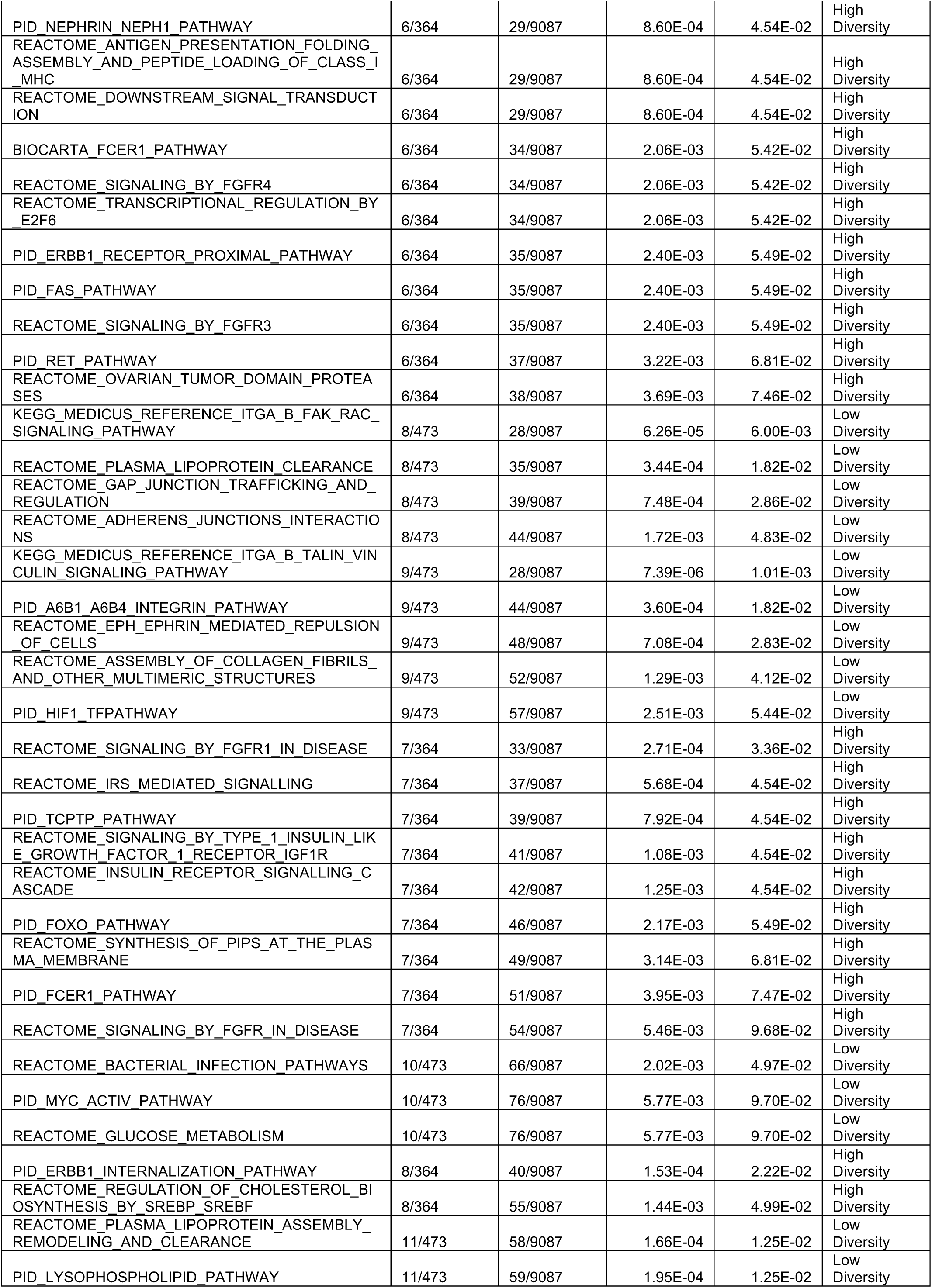
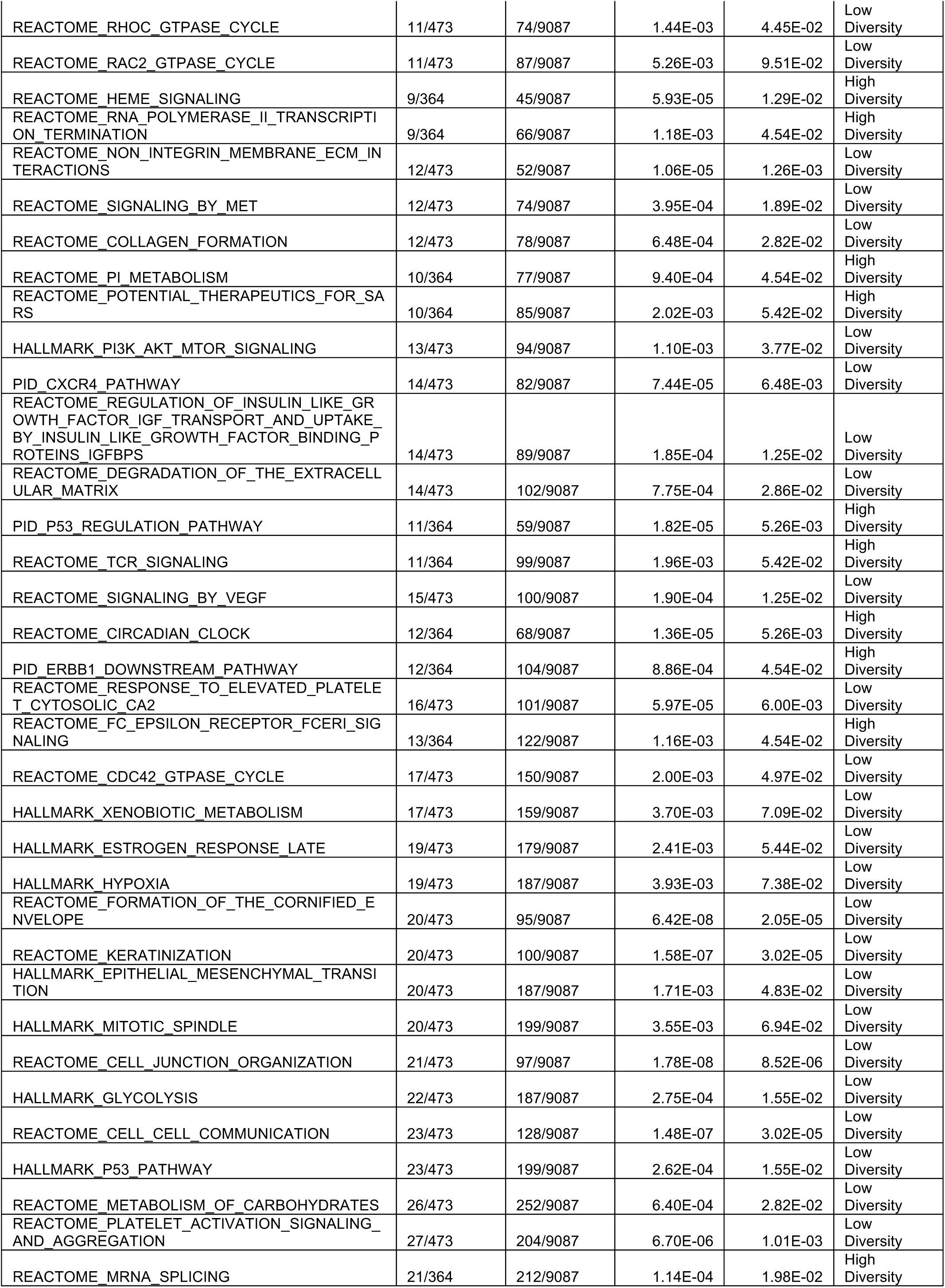
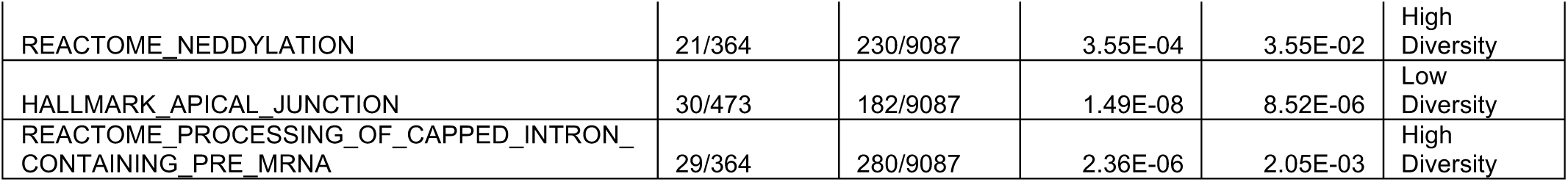
Relevant for Figure 2. Host pathways upregulated in response to low diversity and high diversity microbiomes.

## Notes

### Competing Interest Statement

The authors have declared no competing interest.

